# SyngenicDNA: stealth-based evasion of restriction-modification barriers during bacterial genetic engineering

**DOI:** 10.1101/387985

**Authors:** Christopher D. Johnston, Sean Cotton, Susan R. Rittling, Jacqueline R. Starr, Gary Borisy, Floyd E. Dewhirst, Katherine P. Lemon

## Abstract

Restriction-modification (RM) systems hinder the use of genetic approaches in the vast majority of bacteria. Here, we describe a systematic approach to adapt genetic tools for use in bacteria that are genetically intractable or poorly tractable owing to active RM defenses. In this process, we determine the genome and methylome of a bacterial strain and use this information to define the bacterium’s RM target motifs. We then synonymously eliminate RM targets from the nucleotide sequence of a genetic tool *in silico,* synthesize an RM-silent ‘SyngenicDNA’ tool and propagate the tool as novel minicircle plasmids, termed SyMPL tools, before transformation. Using SyngenicDNA and SyMPL tools, we achieved a profound, >100,000- fold, improvement in the transformation of a clinically relevant USA300 strain of *Staphylococcus aureus* demonstrating the efficacy of these approaches for evading RM systems. The SyngenicDNA and SyMPL approaches are effective, flexible, and should be broadly applicable in microbial genetics. We expect these will facilitate a new era of microbial genetics free of the restraints of restriction-modification barriers.

---

Restriction-modification (RM) systems are bacterial defense mechanisms that have hampered microbial genetic engineering since the inception of recombinant genetics 40 years ago^1^. Found in ~90% of sequenced bacterial genomes, RM systems enable bacteria to distinguish self from nonself DNA via two enzymatic activities: a restriction endonuclease (REase) and a modification methyltransferase (MTase). The REase recognizes the methylation status of DNA at a highly specific DNA target sequence and degrades unmethylated or inappropriately methylated (i.e., nonself) targets. Its cognate MTase protects the same target sequence across the host’s genome via addition of a methyl group, marking each site as self. RM targets vary greatly in sequence and length, typically ranging from 4-18 bp. To date, >450 different target sequences and >4,000 RM-associated enzymes have been identified^2^. Additionally, the number of these systems present in a bacterial cell and the targets recognized are hypervariable and highly species specific, often even strain, specific^3^.

Numerous approaches to overcome RM systems have been developed^4^, almost all involve a mimicry-by-methylation approach to replicate the specific methylation pattern of the desired bacterial host on human-made DNA by using heterologously expressed methyltransferase (MTase) enzymes^5, 6^ or, less successfully, crude cell lysates from the strain of interest. Although sometimes effective, mimicry-by-methylation approaches are time, resource, and cost intensive, and they suffer from limited applicability across different strains (**Supplementary Note 1**).

Here we present SyngenicDNA (**Supplementary Note 2**), a rapid, systematic, and relatively inexpensive approach to circumvent RM barriers during microbial genetic engineering. In contrast to current mimicry-by-methylation approaches, SyngenicDNA involves a stealth-by-engineering approach (**Supplementary Fig. 1**). It is inspired by a simple hypothesis: if a synthetic piece of DNA lacks the highly specific target recognition sequences for a host’s RM systems, then it is invisible to these systems and will not be degraded during artificial transformation. Therefore, in the SyngenicDNA approach, we identify the precise targets of the RM systems within an intractable (or poorly tractable) bacterial strain, eliminate these targets from the DNA sequence template of a genetic tool *in silico*—via single nucleotide polymorphisms (SNPs) or synonymous nucleotide modifications, and synthesize a tailor-made version of the tool that is RM-silent with respect to that specific host (**Fig. 1**). This stealth-based approach allows for simple reworking of currently available genetic tools, and DNA parts, to permit them to efficiently operate in bacteria with active RM defenses. Additionally, for effective propagation of the genetic tool, we have repurposed minicircle technology to eliminate components required in *Escherichia coli* but superfluous in the target host (**Supplementary Fig. 2**). Though not essential, use of minicircle technology reduces the complexity and increases the flexibility of SyngenicDNA.

**Figure 1.**
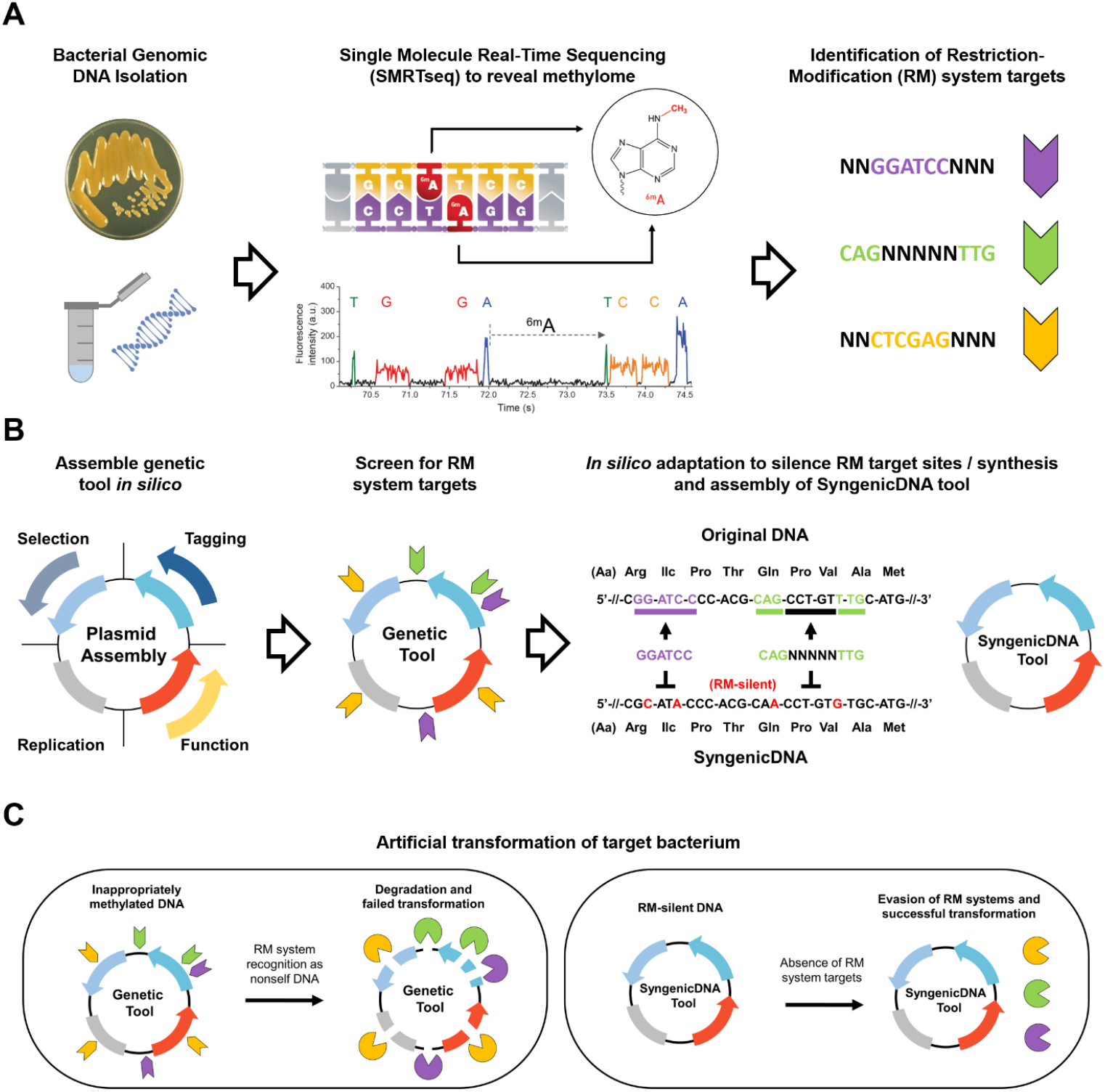
Schematic representation of the SyngenicDNA approach. **(A)** Identification of RM system target motifs by SMRTseq. Methylome analysis of polymerase kinetics during sequencing permits detection of methylated sites at single-nucleotide resolution across the genome, revealing the exact motifs targeted by innate RM systems (indicated by colored nucleotides, where N is any nucleotide) (Kinetic trace image adapted from www.pacb.com). **(B)** Assembly *in silico* of a genetic tool with a desired functionality, followed by screening for the presence of RM target sequences and sequence adaptation, using SNPs or synonymous codon substitutions in coding regions, to create an RM-silent template which is synthetized *de novo* to assemble a SyngenicDNA tool. **(C)** Artificial transformation of the bacterium of interest target bacterium. Inappropriately methylated target motifs of the original genetic tool are recognized as nonself-DNA and degraded by RM systems. In contrast, the SyngenicDNA variant retains the form and functionality of the genetic tool but is uniquely designed at the nucleotide level to evade the RM systems and can operate as desired within the target bacterial host.

There are four basic steps (**Fig. 1**) to produce SyngenicDNA-based genetic tools: 1) target identification, 2) *in silico* tool assembly, 3) *in silico* sequence adaptation, and 4) DNA synthesis and assembly. Below, we detail each step and illustrate the power of the SyngenicDNA method by applying it to a poorly tractable strain of the human pathogen *Staphylococcus aureus* (**Fig. 2**).

**Figure 2.**
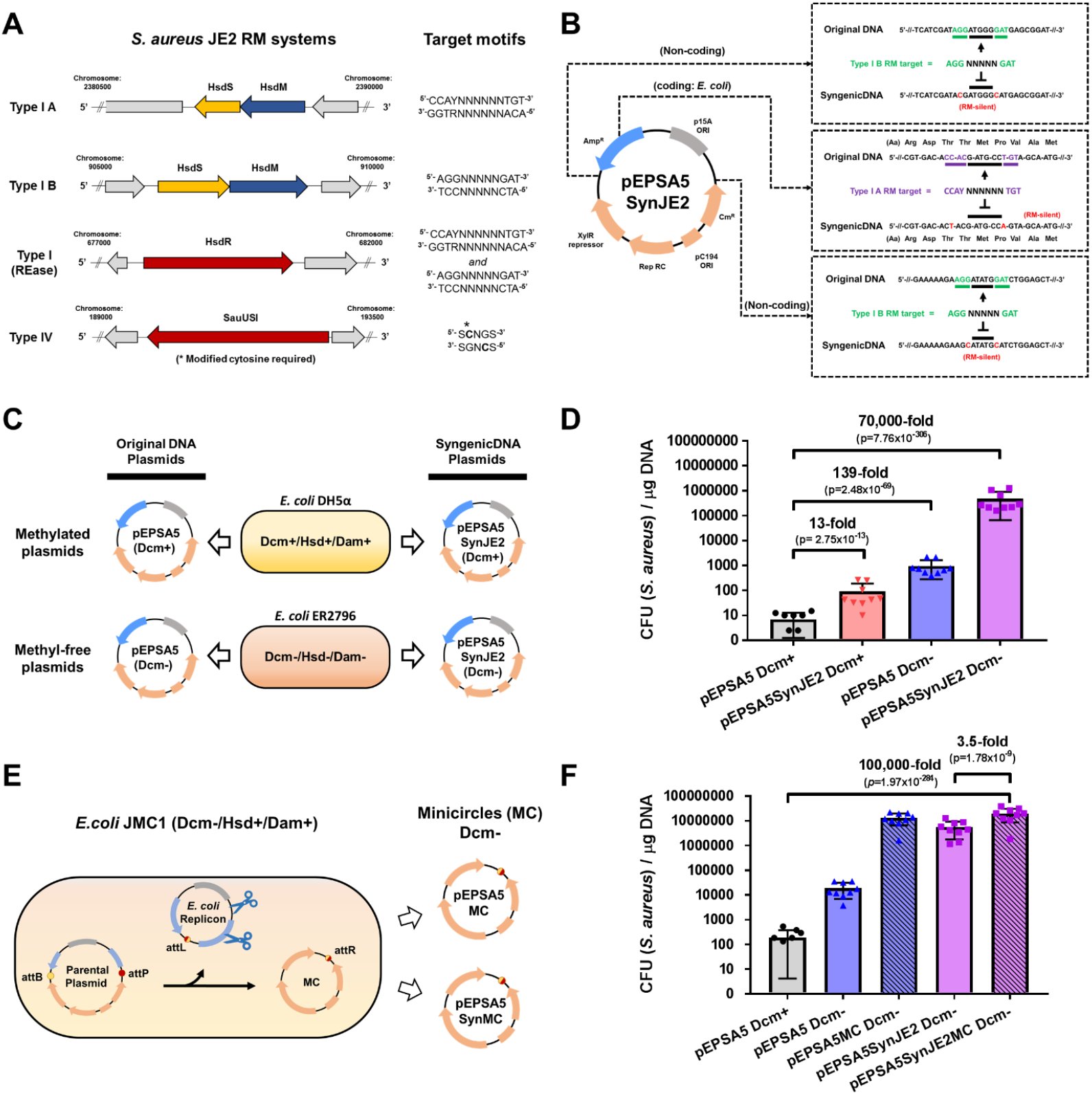
SyngenicDNA and SyMPL approaches applied to *Staphylococcus aureus* JE2. **(A)** JE2 maintains two Type I RM systems and a Type IV restriction system. Restriction endonuclease (HsdR and SauUSI) and methyltransferase (HsdM) genes are shown in red and blue, respectively, and specificity subunit (HsdS) genes are shown in yellow. RM system operons and their corresponding target motifs (where N is any base) were identified by SMRTseq and REBASE analysis. **(B)** Construction of pEPSA5SynJE2, which is an RM-silent variant of the pEPSA5 plasmid tailored to JE2. Six nucleotide substitutions (two synonymous codon substitutions and four SNPs) eliminated all Type I RM system targets from pEPSA5 sequence. **(C)** Plasmid propagation scheme. *E. coli* host strains produce DNA susceptible (DH5α; Dcm+) or resistant *(E. coli* ER2796; Dcm-) to the *S. aureus* JE2 Type IV restriction system. **(D)** Comparison of plasmid transformation efficiency (CFU/μg DNA) with pEPSA5 and the SyngenicDNA-variant pEPSA5SynJE2. **(E)** Propagation of minicircles (pEPSA5MC and pEPSA5SynJE2MC) lacking Dcm-methylated sites within SyMPL producer strain *E. coli* JMC1. (F) Comparison of SyngenicDNA and pEPSA5-based SyMPL plasmid transformation efficiency (CFU/μg DNA) with JE2. Data are means + SEM from nine independent experiments (three biological replicates with three technical replicates each).

Target identification requires the delineation of each methylated site, with single-base resolution, across an entire bacterial genome (i.e., the methylome) and starts with single molecule real-time (SMRT)^7^ genome and methylome sequencing. Using methylome data, we delineate each of the recognition motifs protected by the MTases of the host’s RM systems and infer the targets recognized and degraded by their cognate REases (**Supplementary Note 3**). This yields a concise list of a host microbes’ RM targets to be eliminated from the DNA sequence of the selected genetic tool.

*In silico* tool assembly requires complete annotation of a genetic tool’s sequence with respect to plasmid chassis, replication origins, antibiotic resistance cassettes, promoters, repressors, terminators and functional domains to avoid adverse changes to these structures during subsequent adaptation steps. Ideally, a complete and minimalistic genetic tool with previous demonstrable functionality in a genetically tractable strain is used for initial experiments, allowing for subsequent addition of DNA parts to increase functionality after successful transformation is achieved (**Supplementary Note 4**).

*In silico* sequence adaptation of the genetic tool is the most crucial step of the SyngenicDNA approach and exploits an intrinsic evolutionary weakness present in all RM systems: their exquisite specificity in target sequence recognition. REases are toxic in the absence of their cognate MTases and consequently seldom deviate from their recognition specificity^8^. Accordingly, in this step, we first screen the complete nucleotide sequence of the genetic tool for the presence of RM targets delineated by SMRTseq. We then recode the nucleotides of each RM target *in silico* to eliminate the target while preserving the functionality of the sequence. In noncoding regions, targets are removed by SNPs. In coding regions, the sequence of the target is removed using synonymous codon substitution. A single nucleotide switch is generally sufficient to remove RM targets but multiple switches can also be used. The preferential codon bias of the desired host is used to avoid introducing rare or unfavorable codons during the synonymous switch (**Supplementary Note 5**). Upon complete removal of all RM targets *in silico,* the adapted DNA sequence is RM silent with respect to the host and ready for *de novo* DNA synthesis.

Synthesis and assembly of RM-silent genetic tools is carried out with commercially available *de novo* DNA synthesis and standard assembly approaches, ensuring that any laboratory can construct SyngenicDNA tools. During commercial DNA synthesis, polynucleotide sequences are typically cloned onto an *E. coli* plasmid replicon, which is propagated to yield large amounts of the synthetic DNA. This *E. coli* replicon is convenient but might include RM targets that could lead to degradation of the overall circular tool after transformation into the host species. We have developed two solutions to this potential issue. One solution is to generate a SyngenicDNA *E. coli* plasmid backbone for each specific microbial host strain (**Fig. 2**). However, in routine applications this will increase costs of SyngenicDNA synthesis and, moreover, the *E. coli* replicon itself becomes redundant after propagation in *E. coli,* as it is typically nonfunctional in other bacterial species after transformation. Our alternative solution, therefore, is to remove the *E. coli* replicon entirely using minicircle DNA technology, rather than recode it. This approach also increases flexibility because the same *E. coli* replicon can be used to generate tools for multiple different host strains.

Minicircles (MCs) are minimalistic circular expression cassettes devoid of a plasmid backbone^9^. These are primarily used in gene therapy applications to drive stable expression of transgenes in eukaryotic hosts^10^. MCs are produced by attaching a parental plasmid (PP) to a transgene cassette; cultivating this construct in an *E. coli* host grown to high-cell density; inducing construct recombination to form an isolated transgene on a MC and a separate, automatically degraded, PP containing the *E. coli* replicon; and, finally, purifying isolated MCs by using standard plasmid methods^9^ (**Supplementary Fig. 2**). Because any DNA sequence can take the place of the transgene, we first hypothesized that MC technology could be repurposed to carry entire microbial plasmids and facilitate the removal of superfluous *E. coli* replicons from shuttle vectors. We demonstrated that the incorporation of SyngenicDNA sequences into a PP allowed us to create <u>Sy</u>ngenic DNA <u>m</u>inicircle <u>p</u>lasmid (SyMPL) tools. SyMPL tools include replication, selection, and functional domains for operation in a specific non-E. *coli* host, but lack an *E. coli* replicon despite being isolated at high concentrations from the MC-producing *E. coli* strain. In our SyMPL strategy, we attach a synthesized (and assembled) SyngenicDNA tool to the nonSyngenicDNA *E. coli* PP and propagate this construct in a MC-producing *E. coli* strain. The induction of MCs via recombination, with concurrent induction of a specific endonuclease that eliminates the PP, allows for easy isolation of a minimalistic SyngenicDNA-based genetic tool ready to transform into the desired host strain (**Supplementary Fig. 2**).

Notably, tool propagation in *E. coli* leads to the addition of methylation signatures on plasmid DNA by innate *E. coli* MTase enzymes. We demonstrated that this is also true of the MC-producing *E. coli* host (**Supplementary Fig. 3**). These methylation signatures can activate Type IV restriction systems, which target and degrade methylated DNA motifs, if such systems are present in a desired host strain. Accordingly, to ensure the functionality of our SyMPL approach in a broad range of microbial species, we applied iterative CRISPR-Cas genome engineering^11^ (**Supplementary Fig. 4–6**) to generate a suite of *E. coli* hosts each capable of producing MCs with different methylation signatures, including a methylcytosine-deficient MC producer strain (JMC1: *dcm-, hsdM+, dam+)* (**Supplementary Note 6**).

RM systems are a known critical barrier to genetic engineering in most strains of *Staphylococcus aureus^12^.* Based on its public health importance, we selected *S. aureus* JE2, a derivative of the epidemic USA300 community-associated methicillin-resistant *S. aureus* (MRSA) LAC strain^13^ to demonstrate proof of principle for our stealth-by-engineering approaches. We first applied our SyngenicDNA approach to the *E. coli-S. aureus* shuttle vector pEPSA5 (**Fig. 2A-D, Supplementary Fig. 7**) and generated pEPSA5SynJE2, a variant that differed by only six nucleotides (99.91% identical at nucleotide level), eliminating three RM target motifs present in the original sequence. We demonstrate a ~70,000-fold (*p*=7.76×10^−306^) increase in transformation efficiency (CFU/μg DNA) using the completely RM-silent pEPSA5SynJE2 (propagated in *dcm-E. coli)* compared to the original pEPSA5 plasmid (propagated in *dcm+ E. coli)* (**Supplementary Note 7**).

Subsequently, we sought to determine if a further increase in transformation efficiency could be achieved using the SyMPL minicircle approach. We generated a SyngenicDNA pEPSA5 minicircle for JE2 (pEPSA5SynJE2MC); 38% smaller than pEPSA5 and free of the original *E. coli* replicon (**Supplementary Fig. 8**). This pEPSA5SynJE2MC variant achieved ~2 x 10^7^ transformants/μg DNA, a further 3.5-fold increase (*p*=1.78×10^−9^) in transformation efficiency over pEPSA5SynJE2 and >100,000-fold increase (*p*=1.97×10^−284^) compared to the original unmodified pEPSA5 plasmid (propagated in *dcm+ E. coli).* (**Fig. 2E-F, Supplementary Table 3** and **Supplementary Note 8**). These results demonstrate the profound increase in transformation efficiency that can be achieved by systematic evasion of RM system barriers through stealth-by-engineering approaches.

SyngenicDNA and SyMPL approaches are effective, flexible, and broadly applicable methods to circumvent the RM barriers that heretofore have stymied the advancement of research and development in basic-, synthetic-, and translational-microbiology^14–16^. These methods may also be useful for evasion of other microbial defense mechanisms that rely on distinct target recognition sequences to discriminate self from nonself DNA. The stealth-by-engineering approaches we have developed offer the promise of an era in which microbial genetic system design is unrestrained by a microbe’s innate defense mechanisms.

## METHODS

### Materials

*Escherichia coli* NEBalpha competent cells were purchased from New England Biolabs (NEB) and used as intermediate cloning hosts. *E. coli* ER2796 was provided by the laboratory of Rich Roberts (NEB) and used to produce methylation-free plasmid DNA. *E. coli* MC (ZYCY10P3S2T; original minicircle-producing strain) was purchased from System Biosciences (SBI). Antibiotics and chemicals were purchased from Millipore-Sigma (St. Louis, MO) (Kanamycin, ampicillin, chloramphenicol, spectinomycin, isopropyl-D thiogalactopyranoside; IPTG) or Cayman Chemicals (Anhydrotetracycline). Growth media were purchased from Millipore-Sigma (Luria–Bertani, Brain Heart Infusion) or Oxoid (Vegetable Peptone). DNA isolation kits were purchased from Lucigen (Masterpure Gram Positive kit) and Qiagen (QIAprep Spin Miniprep Kit). Cloning reagents and DNA enzymes were purchased from NEB (Phusion High-Fidelity DNA Polymerase, HiFi DNA Assembly Master Mix, Q5 Site-Directed Mutagenesis Kit, EpiMark Bisulfite Conversion Kit) or Takara (EpiTaq HS for bisulfite-treated DNA). Plasmids were purchased from System Biosciences (SBI) (Parental plasmid; pMC vector), Elitra Pharmaceuticals (pEPSA5), Addgene (pCas; plasmid #42876, pTargetF; #62226) or obtained from the laboratory of George Church, Harvard University (pCKTRBS^1^) or Rich Roberts, NEB (pRRS). Oligonucleotides were purchased from IDT Technologies (Coralville, IA). Electroporation cuvettes (1 mm-gap) were purchased from BioRad and transformations performed on a BioRad Gene Pulser instrument. *De novo* DNA synthesis services and polynucleotide fragments were purchased from Synbio Technologies (Monmouth Junction, NJ). Plasmid DNA sequencing services were purchased from Macrogen (Cambridge, USA) or the DNA core at the Center for Computational and Integrative Biology, Massachusetts General Hospital (Cambridge, MA).

### SMRTseq and RM system identification

The principle of single molecule, real-time sequencing (SMRTseq) and related base modification detection has been detailed previously^2^. SMRTseq was carried out on a PacBioRSII (Pacific Biosciences; Menlo Park, CA, USA) with P6/C4 chemistry at the Johns Hopkins Deep Sequencing & Microarray Core Facility, following standard SMRTbell template preparation protocols for base modification detection (www.pacb.com). Genomic DNA samples were sheared to an average size of 20 kbp via G-tube (Covaris; Woburn, MA, USA), end repaired and ligated to hairpin adapters prior to sequencing. Sequencing reads were processed and mapped to respective reference sequences using the BLASR mapper (https://github.com/PacificBiosciences/blasr) and the Pacific Biosciences’ SMRTAnalysis pipeline (https://www.pacb.com/documentation/smrt-pipe-reference-guide/) using the standard mapping protocol. Interpulse durations were measured and processed for all pulses aligned to each position in the reference sequence. To identify modified positions, we used Pacific Biosciences’ SMRTanalysis v2.3.0 patch 5, which uses an *in silico* kinetic reference and a t-test-based kinetic score detection of modified base positions. Using SMRTseq data, RM system identification was performed essentially as previously described^3^, using the SEQWARE computer resource, a BLAST-based software module in combination with the curated restriction enzyme database (REBASE)^4^. Prediction was supported by sequence similarity, presence, and order of predictive functional motifs, in addition to the known genomic context and characteristics of empirically characterized R-M system genes within REBASE and enabled the reliable assignment of candidate methyltransferase genes to each specificity based on their RM types.

### Bioinformatics and SyngenicDNA adaptation *in silico*

DNA sequence analysis and manipulation was performed using the Seqbuilder and Seqman programs of the DNASTAR software package (DNASTAR, Madison, WI). Codon usage analyses and synonymous substitutions were determined using a combination of CodonW and the Codon Usage Database (www.kazusa.or.jp/codon/), and introduced within Seqbuilder to maintain the amino acid integrity of coding regions within *E. coli.* Clustal Omega (http://www.ebi.ac.uk/Tools/msa/clustalo/) was used to align DNA and amino acid sequences from original ORFs and SyngenicDNA variants. Plasmid DNA (dsDNA) conversions from weight (μg) to molarity (pmol) was performed with Promega BioMath Calculators (https://www.promega.com/a/apps/biomath/).

### DNA synthesis and assembly of SyngenicDNA plasmids

A SyngenicDNA-variant of the pEPSA5 plasmid (pEPSA5Syn) was assembled by replacing a 3.05 kb fragment of the original plasmid, encompassing three JE2 RM target sites, with a *de novo* synthesized DNA fragment that was RM-silent with respect to *S. aureus* JE2 (**Fig. 2, Supplementary Fig.7**). Primers used are listed in **Supplementary Table 1**. The original pEPSA5 plasmid was used as the amplification template for the unmodified backbone, while the plasmid pKan-Frag (Synbio Technologies) was used to amplify the modified RM-silent fragment. PCR amplicons were treated with DpnI to digest non-amplified template DNA and the pEPSA5SynJE2 plasmid was assembled using Gibson cloning. Plasmid nucleotide integrity was confirmed by resequencing. The pEPSA5 and pEPSA5SynJE2 plasmids were propagated within *E. coli* NEBalpha *(dam+, dcm+, hsdM+)* to produce methylated plasmid DNA or *E. coli* ER2796 *(dam-, dcm-*, *hsdM-)* to produce methylation-free plasmid DNA for evasion of Type IV RM systems. Methylation status of plasmid DNA was confirmed by DpnI treatment and agarose gel electrophoresis whereby only methylated plasmids were subject to digestion.

### Constructing an anhydrotetracycline inducible CRISPR-Cas9/λ-Red gene editing system

We utilized a CRISPR-Cas9/λ-Red multigene editing strategy to introduce scarless MTase gene deletions in the *E. coli* MC strain (ZYCY10P3S2T). This strategy, initially described^5^ by Jiang *et al.,* uses a two-plasmid system, pCas and pTarget (**Supplementary Fig. 4A**). In the original system, the pCas plasmid maintains a constitutively expressed cas9 gene and an arabinose-inducible regulatory promoter/repressor module (araC-Pbad) controlling the λ-Red system (Gam, Beta, Exo), both present on a temperature sensitive replicon (repA101Ts). The compatible pTarget plasmid has a sgRNA scaffold for the desired Cas9-target under control of the constitutive promoter (J23119) and a pMB1 origin of replication.

However, since MC formation within the *E. coli* MC strain is also regulated by chromosomally integrated araC-Pbad modules, arabinose induction of λ-Red recombination using the original system would cause unintentional induction of MC-assembly enzymes (the ΦC31 integrase and I-SceI homing endonuclease) during gene editing. To avoid this, we replaced the arabinose-inducible module of the λ-Red system with an alternative tetracycline-inducible module. Primers utilized are listed in **Supplementary Table 1**. A 1318-bp region of pCas, upstream of the λ-Red *gam* gene, containing the araC-Pbad module was replaced with 818-bp tetracycline-inducible regulatory promoter/repressor unit (TetR/PtetO) (**Supplementary Fig. 4B**). The plasmid pCKTRBS served as template DNA for amplification of the TetR/PtetO module, which was spliced to an 11.3 -kb amplicon of pCas (lacking the arabinose module) using Gibson assembly to form pCasTet-λ. The modified pCasTet-λ plasmid, in combination with the original pTarget, allowed for CRISPR-Cas9/λ-Red recombineering using anhydrotetracycline, a derivative of tetracycline that exhibits no antibiotic activity, instead of arabinose as an inducer molecule.

### Genome editing of *E. coli* MC strain

The *E. coli* MC strain contains three active MTases (Dcm+, Hsd+, Dam+) encoded by the *dcm, hsdMS,* and *dam* genes respectively. To create a suite of *E. coli* MC strains, each capable of producing MCs with different methylation signatures, we sequentially deleted (in three-rounds) these MTase genes from the *E. coli* MC genome using our modified anhydrotetracycline-inducible CRISPR-Cas9/λ-Red recombineering strategy (**Supplementary Fig. 4–6**). In this strategy, λ-Red mediated recombination with a DNA editing template eliminates the MTase gene from the chromosome, followed by CRISPR-Cas9 mediated targeting of the MTase gene in unedited cells. Double-stranded DNA breaks introduced by CRISPR/Cas9 are toxic in bacteria, so only cells for which the target sequences have been edited can survive, allowing for positive selection of recombination events. MTase deletion template plasmids were constructed by assembling PCR amplicons of regions 5’ and 3’ of each MTase (reflecting the desired deletion event) onto a pRRS plasmid backbone (**Supplementary Fig. 4C**). These pRRS-based template plasmids were then used to PCR amplify linear editing templates for λ-Red recombineering. To remove template plasmid-carryover during electrotransformation, editing template amplicons were DpnI treated and PCR purified prior to use.

*E. coli* MC competent cells (System Biosciences) were first transformed with pCasTet-λ to form *E. coli* JMC, which constitutively expressed the Cas9 protein but lacked a gRNA target (**Supplementary Fig. 5**). JMC electrocompetent cells (harboring pCasTet-λ) were generated as previously described^6^. For λ-Red induction of JMC cells, anhydrotetracycline (200 ng/ml; ~0.5 μM) was added to the growing (30°C) culture 30 min prior to making cells competent, as described for the arabinose-based system^6^.

In the first round of genome editing, electrocompetent JMC cells were transformed with the *dcm*-deletion editing template and pT-Dcm (pTarget with a single gRNA targeting the *dcm* gene, under control of the J23119 constitutive promoter). For electroporation, 50 μl of cells were mixed with a 5 μl combination of 100 ng pT-Dcm plasmid and 200ng dcm-deletion editing template DNA; electroporation was performed in a 2-mm Gene Pulser cuvette (Bio-Rad) at 2.5 kV. Cells were recovered at 30°C for 1 h before selective plating at 30°C on LB agar containing kanamycin (50 μg/ml) and spectinomycin (50 μg/ml). Transformants were identified by colony PCR and DNA sequencing. Primers are listed in **Supplementary Table 1**. After confirmation of *dcm* deletion, the edited colony harboring both pCasTet-λ and pT-Dcm was cured of the latter plasmid by IPTG induction (0.5 mM), essentially as described previously^5^. Briefly, IPTG induces the production of gRNA, which targets the origin of replication of pT-Dcm after interaction with the constitutively expressed Cas9 protein. This gRNA is encoded on the pCasTet-λ plasmid under transcriptional control of the lacO/LacI (IPTG-inducible) system. The resulting *E. coli* strain, (dcmΔ/pCasTet-λ+) was made competent once again for the next round of editing, or cured of the pCasTet-λ plasmid by incubation at 37°C for four continuous inoculums, to form a plasmid-free minicircle producing strain *E. coli* JMC1 *(dcm-, hsdM+, dam+).*

In the second round of genome editing, the entire process was repeated targeting the Hsd MTase system. *E. coli* dcmΔ/pCasTet-λ+ was transformed with the Hsd-deletion editing template and the pT-Hsd plasmid (pTarget with a single gRNA targeting the *hsdM* gene). The resulting *E. coli* strain, (dcmΔ,hsdMΔ, pCasTet-λ+) was cured of the pCasTet-λ plasmid to form the *E. coli* JMC2 strain *(dcm-, hsdM-, dam+).* In the third round, the entire process was repeated targeting the Dam MTase system. *E. coli dcm-, hsdM-,* pCasTet-λ+ was transformed with the dam-deletion editing template and the pT-Dam plasmid (pTarget with a single gRNA targeting the *dam* gene). The resulting *E. coli* strain *(dcm-, hsdM-, dam-)* was cured of both plasmids to form the completely methyl-free *E. coli* JMC3 strain *(dcm-, hsdM-,* dam-).

After each round of genome editing, the phenotypic effect of *dcm, hsdM,* and *dam* gene deletions were confirmed using bisulfite sequencing, SMRTseq, and methyl-dependent restriction enzyme analysis, respectively (**Supplementary Fig. 6**). Site directed bisulfite sequencing and DpnI methyl-dependent restriction analysis of gDNA were performed essentially as we described previously^7^.

### Production of SyMPL tools

The 4.3 kbp *S. aureus* replicon of both pEPSA5 plasmids (pEPSA5 and the pEPSA5SynJE2) were PCR amplified and spliced to the MC parental plasmid (pMC; Systems Biosciences) to form pEPSA5P and pEPSA5SynJE2P (P denotes parental). Primers listed in **Supplementary Table 1**. To evade the Type IV restriction system of *S. aureus* JE2, which targets Dcm-methylated cytosine residues, we used our *dcm*-deficient MC-producing *E. coli* strain JMC1 *(dcm-, hsdM+, dam+).* Competent plasmid-free *E. coli* JMC1 cells, prepared as described previously, were transformed with pEPSA5P and pEPSA5SynP. Minicircle induction and isolation was performed per manufacturers recommendations for the original *E. coli* MC strain (ZYCY10P3S2T). The resulting SyMPL tools pEPSA5MC and pEPSA5SynMC were eluted in high pure H_2_0 and normalized to 250 ng/μl prior to transformation. Plasmid nucleotide integrity was confirmed by resequencing.

### *Staphylococcus aureus* transformations

Electrocompetent *S. aureus* JE2 cells were prepared using a modified version of that used by Löfblom *et al.*^8^ Briefly, overnight cultures of *S. aureus* JE2 (~OD600nm=1.8) in vegetable peptone broth (VPB) were diluted to an OD600nm of 0.25 in fresh prewarmed VPB. In initial experiments to test the efficacy of the SyngenicDNA method, cultures were grown at 37°C with shaking (100 rpm) until they reached an OD600nm between 0.8-0.95 (~3 hours). However, in the interim of SyngenicDNA experiments and SyMPL method experiments, we observed increased JE2 cell competency was achieved when cultures were grown to an OD600nm between 1.5-1.7 (~6 hours). Therefore, we performed all SyMPL experiments with cells harvested at this higher optical density. In both cases, when culture tubes reached the desired OD, culture flasks were chilled on wet ice for 15 min. Cells were harvested by centrifugation at 5000 x g at 4°C for 10 min, washed once in equal volumes of ice-cold sterile water and pelleted at 4°C. The cells were then washed in 1/10 volume ice-cold sterile 10% glycerol, repeated with 1/25 volume ice-cold sterile 10% glycerol, repeated with 1/100 volume ice-cold sterile 10% glycerol, resuspended in 1/160 volume of ice-cold sterile 10% glycerol and then aliquoted (250 μl) into 1.5 ml tubes. Electrocompetent cell aliquots were frozen at −80°C until use.

For electroporation, a single aliquot was utilized for each individual experiment for accurate comparison of transformation efficiency between plasmids. The aliquot was thawed on ice for 5 min, transferred to room temperature for 5 min, centrifuged at 5000 x g for 1 min and resuspended in 250 μl sterile electroporation buffer (10% glycerol, 500 mM sucrose). A 50 μl volume of competent cells was mixed with 1 μg plasmid DNA (250 ng/ul in sterile water) and added to a sterile 1mm-gap electroporation cuvette. The cells were pulsed once using a Bio-Rad Gene Pulser System (settings: 25 μF, 100 Ω, 2.1 kV with a 2.3 millisec time constant) and outgrown in 1 ml of trypic soy broth with 500 mM sucrose for 1 hour at 37°C, diluted for spreading on trypic soy agar plates with 15 μg/ml Cm and incubated overnight at 37°C.

### Scientific Rigor and Experimental Design

Transformation efficiencies (presented in Figure 2 D and F) were determined based upon nine independent experiments. We prepared three independent batches of electrocompetent *S. aureus* cells (Biological Replicate 1,2, and 3; **Supplementary Table 3**). Three aliquots from each batch of electrocompetent cells were used to perform three independent transformation experiments, typically on consecutive days (Technical Replicates A, B, and C; **Supplementary Table 3**). A single plasmid preparation (for each pEPSA5 variant) was used for all technical replicates within a batch. A fresh plasmid preparation (for all pEPSA5 variants) was used for each new batch of cells to account for variation associated with plasmid propagation/isolation from *E. coli* strains and the effect of freeze-thaw on plasmid DNA. In independent experiments, a single 250 μl aliquot of electrocompetent *S. aureus* was used for all plasmids (50 μl/plasmid) within each of the nine experiments, so that data within technical replicates could be treated as paired, or “clustered” across the four plasmids, and plasmid transformation efficiencies could be compared validly and efficiently. The average of CFU counts from a minimum of three replicate agar plates was used when determining transformation efficiencies for individual plasmids within experiments.

### Statistical analysis

Statistical analyses were carried out using Graphpad Prism (version 7.04; GraphPad Software, San Diego, CA) and Stata version 12.1 (StataCorp. 2011. Stata Statistical Software: Release 12. College Station, TX: StataCorp LP). Means with standard error (SEM) are presented in each graph. As appropriate for count data, we compared transformation efficiency across plasmids by fitting negative binomial regression models with two-sided alpha=0.05. We used a generalized estimating equations (GEE) framework and robust standard errors to account for clustering within technical replicates of competent cells. For each experiment designed as a 2×2 factorial design, we fit main effects and multiplicative interaction terms (see Experimental Design). This can be thought of as a difference-in-differences analysis, quantifying how the effect of one condition (e.g. SyngenicDNA plasmid versus unmodified plasmid) differs in the presence or absence of another condition (e.g. propagated in a Dcm+ or a Dcm-*E. coli* host).

### Data availability

Complete genome sequences and associated methylome annotations of *Staphylococcus aureus* USA300 JE2_Forsyth and *Escherichia coli* MC_Forsyth have been submitted to REBASE (http://rebase.neb.com/) for public release under organism # 21742 and # 21741, respectively. The nucleotide sequences of each plasmid used in this study are included here as Supplementary files. Raw CFU colony count data for determination of transformation efficiencies, along with data for associated analyses, are presented in Supplementary Tables 3 – 5.

## SUPPLEMENTARY NOTES

### Supplementary Note 1

Numerous approaches have been developed in attempts to overcome the restriction-modification (RM) barriers and each of these provides evidence that circumvention of RM systems can lead to genetic tractability in bacteria^1–8^. These can all be referred to as mimicry-by-methylation approaches, as they essentially seek to modify the methylation pattern of a genetic tool to match the desired host and achieve molecular mimicry. Such approaches can be categorized as either unrefined or sophisticated, and each has a distinct disadvantage^9^, including being arduous, resource intensive, or requiring many years to develop. Once developed, these typically suffer from limited applicability across different strains^10^.

There are three common unrefined approaches. **(1)** Randomly mutagenize entire populations of a genetically intractable bacterial strain with ultraviolet radiation or chemicals, and then select for restriction-deficient mutants with increased transformation efficiency^11, 12^. Although random mutagenesis may generate genetically tractable mutants, it also generates additional undefined mutations, such that genetically tractable mutants are no longer a good model of the original strain^13^. **(2)** Mix plasmid DNA with a crude cell extract from the strain of interest so that the strains’ innate MTases methylate (protect) targets present on the plasmid, thus marking them as self DNA^14^. **(3)** Expose the target bacterium to a nonlethal heat treatment before transformation to temporarily inhibit restriction enzyme activity^15, 16^. Neither heat-nor crude extract-treatments are reproducible, and both have limited effectiveness among most strains^3, 16^.

There are two common sophisticated approaches. **(1)** Methylate target sites on tools by using *in vitro* methylation with either recombinant MTases^14, 17^ or commercially available MTase enzymes^18^, which are currently available for only 37 of >450 known targets^19^. **(2)** Alternatively, achieve *in vivo* methylation by passaging a plasmid through a related strain that is either restriction enzyme deficient^20^ or engineered to generate the methylation profile of the strain of interest, i.e., the plasmid artificial modification (PAM) technique^21^. Although the PAM approach is useful in some strains, inherent complications prevent its widespread application. These include; **a**) the exact genetic loci and accessory open reading frames (ORFs) for most RM systems are poorly defined and therefore cannot be introduced into *E. coli* PAM hosts without extensive trial and error^22^; **b**) many MTases cannot be cloned functionally in the absence of their REase partner or are difficult to clone in *E. coli* due to differences in promoter structure, GC content, codon usage, or host-toxicity; and c) many bacteria have multiple active RM systems and multilayered methylated signatures that become difficult to effectively recapitulate *in vitro,* i.e., it is impractical to clone several functional MTase enzymes within a single *E. coli* PAM host for each new bacterial strain of interest.

### Supplementary Note 2

SyngenicDNA is an intentionally broad term. The word syngenic (also syngeneic) is used in immunology to mean sufficiently identical and immunologically compatible as to allow for transplantation without provoking an immune response. It is usually applied in the context of eukaryotic cell or tissue transplantation in cases where the donor and the recipient are identical twins^23^. Syngenic transplants are the easiest because the identical recipient’s immune system readily accepts the graft without risk of rejection. Accordingly, we coined the term ‘SyngenicDNA’ and define it as a piece of synthetic DNA that has been engineered with sufficient sequence and epigenetic compatibility to allow it to function as self within a specific bacterial host, upon artificial transformation, and be accepted by its RM defenses, i.e., the bacterial innate immune system^24^.

### Supplementary Note 3

Post-replicative modification of DNA by MTases in bacteria results in three types of epigenetic markers: N6-methyladenine (m6A), N4-methylcytosine (m4C), and 5-methylcytosine (m5C)^22^. The complete set of methylations across a bacterial genome is termed the methylome. Currently, efficient methylome analysis can only be accomplished by using single molecule real-time sequencing (SMRTseq; www.pacb.com)^25^. During SMRTseq, a polymerase adds fluorescently labelled bases to a DNA template while the sequencing instrument records both the sequence of bases added and the kinetic information (milliseconds) between successive additions, forming a sequencing trace. DNA templates containing a methylated base cause the polymerase to stall at those sites, leading to a delay in the sequence trace. This kinetic information is used to identify the specific sites of methylation in genomic DNA (m6A, m4C or m5C) based on their characteristic trace^25^. SMRTseq analysis software summarizes the exact sequence of the methylated motifs, the number of motifs present on the genome and the percentage of motifs that are methylated.

Accordingly, during the target identification step within the SyngenicDNA approach, we use SMRTseq-generated methylome data to identify active RM systems and then infer the specific target recognized by the REase of each system. In a bacterial genome, a methylated motif represents either **A**) an RM system’s target recognition sequence methylated by an MTase to protect the site from its cognate REase, or **B**) a modification introduced by an orphan MTase, which lacks a cognate REase and may be involved in regulatory activity^26^. To differentiate between these two possibilities, we first evaluate the quantitative SMRTseq methylome data. An active RM system methylates 100% of its target motifs in the genome, because unmethylated motifs are substrates for the cognate REase, which introduces chromosomal breaks resulting in bacterial cell death^27, 28^. Therefore, allowing for a small margin of incomplete post-replicative methylation in actively dividing cells during DNA isolation, we assume that motifs that are methylated >95% indicate an active RM system (**Supplementary Fig. 3A**). Second, we use REBASE analysis (Methods) to confirm suspected orphan MTases as described previously^19^. We designate MTases as orphans if we cannot detect an REase gene homolog with the same target site less than 10 genes away from the MTase, based on genomic coordinates^22, 29^. Thus, upon completion of the target identification step, we have a concise list of the target sequences of a strain’s active RM systems, targets that need to be eliminated from the DNA sequence of the selected genetic tool *in silico.*

### Supplementary Note 4

The SyngenicDNA approach is most readily applicable to genetic tools that are functional in tractable bacterial strains, to modify them for use in strains that are currently intractable or poorly tractable due to RM barriers (**Figure 2**). In addition, the SyngenicDNA approach can facilitate modular assembly of new genetic tools for synthetic biology applications.

Synthetic biology focuses on the construction of biological parts that can be understood, designed and tuned to meet specific criteria^30^, with the underlying principle that genetic tools should be minimalistic, constructed of modularized parts and sequence optimized to allow for compatibility. Standardized formats for genetic tool assembly exist to facilitate the simple implementation of synthetic tools and the distribution of physical parts between different laboratories^30–33^; with the BioBrick standard being the most commonly adopted^32^. However, because RM systems vary between different strains of the same bacterial species^22^, the design of reusable DNA parts that require physical assembly for different bacteria is generally not applicable for intractable or poorly tractable strains with active RM systems.

In the SyngenicDNA approach, we adopt the core principles of synthetic biology, modularity and compatibility, and account for variation in bacterial RM systems between strains by removing the need for physical assembly of re-used parts propagated in other bacterial species. Because SyngenicDNA-based genetic tools require DNA synthesis *de novo* in the later step, the *in silico* tool assembly step can be utilized to augment plasmid backbones with additional useful parts (e.g., antibiotic resistance cassettes, promoters, repressors, terminators and functional domains, such as transposons or fluorescent markers) or create new tools. Additionally, because there is no requirement for a laboratory to physically obtain template DNA for PCR amplification of these additional parts, researchers only need access the publicly available DNA sequences of new parts to integrate them into a SyngenicDNA-based genetic tool, which are then synthetized *de novo* in context. Compatible replication origins and accessory elements for the majority of cultivable bacterial phyla can be obtained from **A**) the NCBI Plasmid Genome database, containing >4418 complete DNA sequences of bacterial plasmids^34^ **B**) the European Bioinformatics Institute (EMBL-EBI) Nucleotide Archive, which maintains >1000 plasmid sequences (https://www.ebi.ac.uk/genomes/plasmid.html) or **C**) the ACLAME database (A CLAssification of Mobile genetic Elements), which maintains an extensive collection of mobile genetic elements (MGEs) including microbial plasmids from various sources (http://aclame.ulb.ac.be/).

### Supplementary Note 5

The frequency with which an RM target occurs in the DNA sequence of a genetic tool depends on the length and base composition (GC vs AT content) of the target motif. Target motifs vary greatly in sequence and length, ranging from 4-18 base pairs (bp), with >450 different motifs identified to date^19^. RM systems are classified into four types (Type I, II, III, and IV), based on their target motifs recognized and, also, their subunit composition, cleavage position, cofactor requirements, and substrate specificity^24^. Type I-III systems, with exceptions, recognize and cut a target sequence if it lacks an appropriate methyl group. Characteristically, Type I systems target discontinuous bipartite DNA motifs comprising two specific half-sequences separated by a nonspecific spacer gap of 6 to 8 bp. One of the best characterized examples is the EcoKI system that recognizes AACN6GTGC (N is any base)^35^. Type II systems are a conglomeration of many different subsystems that target both continuous and noncontinuous motifs ranging from 4 bp (e.g., AGCT of the AluI system^36^) to 15 bp (e.g., CCAN9TGG of the XcmI system^37^). Type III systems recognize short continuous asymmetric targets ranging from 4 bp (e.g., CGCC of the TmeBIV system^19^) to 7 bp (e.g., AGCCGCC of the Bpe137I system^19^). Type I-III RM system targets that occur within non-coding regions can be eliminated readily using single nucleotide polymorphisms (SNPs), whereas those that occur in coding regions require synonymous codon switches (**Fig. 2B**).

Many genetic tools are dual host-range plasmids (i.e., shuttle vectors) composed of two different functional replicons (origin of replication and accessory genes) permitting them to operate in multiple bacterial species (usually a laboratory strain of *E. coli* and another desired host species). The activity of the two replicons is usually partitioned depending on the bacterial host strain. The *E. coli* replicon is active when propagating the genetic tool in *E. coli* while the other replicon remains inactive until transferred to the desired host strain, whereupon the *E. coli* replicon then becomes inactive.

Notably, bacteria use synonymous codons at unequal frequencies, with some favored over others by natural selection for translation efficiency and accuracy, known as codon bias^38^. Therefore, to avoid the introduction of rare or unfavorable codons when eliminating RM targets within a genetic tool *in silico,* it is critical to distinguish on which replicon each target motif is present and introduce synonymous substitutions corresponding to the codon bias of that specific host. Codon bias is determined by annotation and analysis of the host’s genome generated by SMRTseq. For example, the pEPSA5 plasmid^39^ is an *E. coli–S. aureus* shuttle vector containing a 2.5 kb *E. coli* replicon (ampicillin-resistance gene and low copy number p15a origin for autonomous replication) and a 4.3 kb *S. aureus* replicon (chloramphenicol-resistance gene, pC194-derived origin, and a xylose repressor protein gene, *xylR)* (**Supplementary Fig. 2B**). The *S. aureus* replicon is nonfunctional when pEPSA5 is maintained and propagated within *E. coli,* and vice versa. Therefore, RM targets that occurred within a coding region of the pEPSA5 *E. coli* replicon were modified with synonymous substitutions adhering to *E. coli* codon bias (**Methods)**.

Additionally, if an RM target motif identified corresponds to a commercially available MTase enzyme, one could utilize *in vitro* methylation of SyngenicDNA-based tools (downstream of *de novo* synthesis) rather than elimination of such targets via nucleotide substitution. This will decrease the total number of necessary substitutions and reduce the likelihood of introducing unfavorable alterations. However, of the >450 motifs identified to date, only 37 of these targets are represented by available MTase enzymes. Furthermore, only 16 of those available commercially are isolated MTase enzymes that are useful for *in vitro* DNA methylation (**Supplementary Table 2**): the remaining 21 enzymes exist as RM complexes, with MTase and REase subunits that compete for enzymatic modification and restriction activities, respectively^19^. Nevertheless, in cases where an MTase is available, all other RM targets could be eliminated *in silico* to generate stealth-by-engineering and a foundational SyngenicDNA genetic tool synthesized and assembled, followed by *in vitro* methylation prior to transformation.

In contrast to Type I-III systems detailed above, Type IV restriction systems lack MTases and instead are composed of methyl-dependent REase enzymes that only cleave DNA sequences with methylated, hydroxymethylated, or glucosyl-hydroxymethylated bases within their short target motifs. These systems are exemplified by the *Staphylococcus aureus* system SauUSI^40^; a modified cytosine restriction system targeting S^5m^CNGS (either^m5^C or^5hm^C) where S is C or G. The presence of such systems in a bacterial host have significant implications for genetic engineering due to their repressive effect on transformation efficiency (**Fig. 2D**). It is relatively simple to detect the presence of a Type IV system in a genome by screening for homologs to the >8,210 putative Type IV REases in REBASE^19^. However, identification of Type IV system target motifs is inherently more difficult than for Type I-III systems because their targets motifs cannot be determined through SMRTseq and methylome analysis owing to the absence of an indicative epigenetic modification on host gDNA^22^. Neverthless, the unintentional activation of Type IV systems can be avoided by the propagation of SyngenicDNA based tools in an intermediate E. *coli* host that does not methylate DNA *(dam-, dcm-,* hsdRMS-)^41^, thus avoiding recognition and degradation by any Type IV systems present.

As such, the systematic identification of the specific RM barriers present within a bacterial host facilitates the development of a tailored stealth-by-engineering stratagem to evade these barriers during genetic engineering. Once developed, this stratagem can then be reapplied to create additional SyngenicDNA based genetic tools for the same host strain.

### Supplementary Note 6

The majority of laboratory *E. coli* strains, including the MC producing *E. coli* host used in this study, contain three active MTases (Dam, Dcm, and HsdM) that introduce methylation modifications to specific target sites on the host genome (**Supplementary Fig. 3**). The Dam MTase modifies the adenine residue (^m6^A) within the sequence GATC, the Dcm MTase modifies the internal cytosine residue (^m5^C) of the sequence CCWGG (where W is A or T), and the HsdM MTase modifies the internal adenine residue (^m6^A) of the sequence AACN6GTGC. Therefore, plasmid tools propagated within such *E. coli* strains, including the minicircle (MC)-producing strain (ZYCY10P3S2T), are modified at these targets sequences.

The presence of methylated sites on SyngenicDNA-based tools could activate Type IV RM systems upon artificial transformation. Generally, unintentional activation of Type IV systems is avoided by the propagation of plasmids within methyl-deficient *E. coli* strains such as JM110 *(dam-, dcm-, hsdRMS+)* or ER2796 *(dam-, dcm-, hsdRMS-),* thus preventing recognition and degradation via these systems. However, such methyl-free *E. coli* strains are unable to produce MCs since construction of the *E. coli* MC-producing strain^42^ required complex engineering to stably expresses a set of inducible minicircle-assembly enzymes (the øC31-integrase and the I-SceI homing-endonuclease for induction of MC formation and degradation of the parental plasmid replicon, respectively). Accordingly, when we repurposed MC technology for bacterial applications, it was also necessary to engineer *E. coli* MC producer strains that generates various forms of methylation-free MCs.

Although a completely methylation-free MC producer could be required when working against Type IV systems targeting both adenine- and cytosine-methylated DNA, bacterial RM systems exist with targets that specifically match the *E. coli* Dam MTase motif (GATC), such as the Dpn system of *Streptococcus pneumoniae^43^* or the Pin25611FII system of *Prevotella intermedia^22^.* These systems digest unmethylated Dam sites on genetic tools propagated within a completely methyl-free strain, hence Dam methylation is protective in these cases. Therefore, we created a suite of *E. coli* strains capable of producing distinct types of methyl-free MC DNA (**Supplementary Fig. 4-6**) to account for the inherent variation of RM systems in bacteria and maximize the applicability of our SyMPL approach,

We applied iterative CRISPR-Cas9 genome editing to sequentially delete MTase genes from the original *E. coli* MC producer strain *(dam+, dcm+, hsdM+)* (**Supplementary Fig. 4–6**). These new strains produce methylcytosine-free MC DNA *(E. coli* JMC1; *dam+, dcm-, hsdM+),* methylcytosine- and methyladenine-free MC DNA except for Dam methylation *(E. coli* JMC2; *dam+, dcm-, hsdM-),* and completely methyl-free MC DNA *(E. coli* JMC3; *dam-, dcm-, hsdM-).* Depending upon the Type IV RM systems identified within a desired bacterial host, one of these strains can be selected and utilized for production of SyMPL tools.

### Supplementary Note 7

By definition, an entirely SyngenicDNA plasmid is silent with respect to all (Type I, II, III, and IV) RM systems within a host strain and is designed to maximize transformation efficiency. In addition, generation of complementary sets of partially SyngenicDNA plasmids can be used to determine the relative contribution of different RM systems within a host strain. For example, *S. aureus* JE2 contains two active Type I RM systems, which target unmethylated bipartite sequence motifs, in addition to a Type IV restriction system, SauUSI^40^, that targets methylated S^5m^CNGS motifs (either ^m5^C or ^5hm^C) where S is C or G (**Fig. 2A**). Plasmid tools propagated in *E. coli* strains containing the Dcm orphan MTase are methylated at C^5m^CWGG motifs, which overlap with the SauUSI target motif resulting in vulnerability to degradation by this restriction system upon transformation to *S. aureus.* Therefore, in addition to the fully SyngenicDNA plasmid (pEPSA5SynJE2Dcm-) we generated partially SyngenicDNA plasmids, one that is RM-silent to Type I systems but not to Type IV systems (pEPSA5SynJE2Dcm+) and another that is vice versa (pEPSA5Dcm-) to determine the relative contribution of Type I or Type IV systems to the genetic barrier in *S. aureus* JE2. This type of experimental approach can be viewed as a 2×2 factorial design, crossing silencing of the Type I systems and silencing of the Type IV system.

The original pEPSA5 plasmid propagated in *E. coli* NEBalpha, a standard Dcm+ laboratory strain, achieved consistently poor transformation efficiencies in our hands (~10 CFU/μg DNA). This plasmid contains 11 individual RM target motifs (Type I; n=3, and Type IV; n=8, **Supplementary Fig. 7A**). Both system types are known to be actively involved in defense from foreign DNA in *S. aureus^1^’*^44–46^. Elimination of only Type I target motifs from the plasmid (pEPSA5SynJE2Dcm+) achieved a 13-fold increase (*p*= 2.75×10^−13^) in transformation efficiency. In contrast, elimination of only Type IV system targets, by passaging pEPSA5 through the Dcm-deficient strain *E. coli* ER2796 (pEPSA5Dcm-), achieved a >139-fold increase (*p*=2.48×10^−69^) in efficiency. However, when both Type I and Type IV targets were eliminated (pEPSA5SynJE2Dcm-), we observed a supra-multiplicative (rather than an additive) effect on transformation efficiency, with in an increase of ~70,000-fold (*p*=7.76×10^−306^) compared with the original pEPSA5Dcm+ plasmid (p for interaction=6.98×10^−27^). The mechanism of this supra-multiplicative effect is not immediately apparent and raises questions for future studies. For example, are there direct interactions between the distinct types of systems? Additionally, comparing the original and the SyngenicDNA pEPSA5 plasmids independently of the Type IV system (pEPSA5Dcm-and pEPSA5SynJE2Dcm-) showed that elimination of the three Type I system targets achieved ~500-fold increase (*p*=<1.0×10^−306^) in efficiency (**Fig. 2B**). As demonstrated here, by using system-specific sets of partially RM-silent plasmids, we can detect the relative contributions of different RM systems within a host strain.

### Supplementary Note 8

We used the *dcm-* strains *E. coli* ER2796 and *E. coli* JMC1 to carry out the minicircle (MC) experiments independently of the Type IV system in *S. aureus* JE2. The majority of the *S. aureus* JE2 RM system targets present on pEPSA5 are in the *E. coli* replicon (Type I; n=2, and Type IV; n=8) with only a single Type I system target in the *S. aureus* replicon (**Supplementary Fig. 7A**), thus the MC approach eliminates 2 of the 3 Type I targets. The focus here was on investigating 1) whether the SyMPL approach achieves equal or perhaps even greater efficiency than the SyngenicDNA approach, 2) whether removal of all Type I targets is required to achieve appreciable gains in transformation efficiency (compared with a partially SyngenicDNA plasmid that has a single Type I target remaining), and 3) whether removing all Type I targets adds further gains in transformation efficiency compared with leaving a single Type I target intact. The original plasmid pEPSA5 (Dcm+) was included in experiments only as a control for accurate final comparison of efficiencies and was not considered a primary comparison.

Importantly, in SyMPL experiments, by reducing the overall size of MC plasmids, we also increased the number of *S. aureus* replicons present within the μg of DNA used for transformations (compared with the μg used for full-length plasmids). Increasing the yield of functional replicons/μg of DNA might be an additional advantage of the MC approach. Thus, to more accurately compare transformation efficiencies between MCs and full-length plasmids, we performed a secondary analysis in which we adjusted the transformation efficiencies from CFU/μg DNA to CFU/pmol DNA (**Supplementary Table 5, Supplementary Fig. 9**).

On a CFU/pmol DNA basis, the MC variant pEPSA5MCDcm-achieved a 436-fold increase in transformation efficiency over the original plasmid pEPSA5Dcm-(*p*=<1.0×10^−306^). This increase could be due to the elimination of the two Type I target motifs along with the *E. coli* replicon in the MC variant (**Supplementary Fig. 8**), the smaller MCs passing more readily through the reversible pores formed in the *S.aureus* cell envelope during electroporation, or a combination of both. The mere 2.3-fold (*p*=1.29×10^−4^) increase in transformation efficiency achieved by MC variant pEPSA5SynJE2MC over the plasmid pEPSA5SynJE2, both of which are completely RM-silent in JE2, favors the first possibility.

In contrast, pEPSA5MC and pEPSA5SynJE2MC differed only by the presence or absence of a single Type Itarget, respectively (**Supplementary Fig. 7A**). Eliminating this single target sequence resulted in a modest 1.5-fold (*p*=1.01^−14^) increase in transformation efficiency. This suggests that in future applications of the SyngenicDNA approach, if a single target exists in an unadaptable region of DNA, such as an origin of replication or a promoter, its inclusion on an otherwise RM-silent plasmid might have minimal impact on the overall transformation efficiency.

## SUPPLEMENTARY FIGURES

**Supplementary Figure 1.**
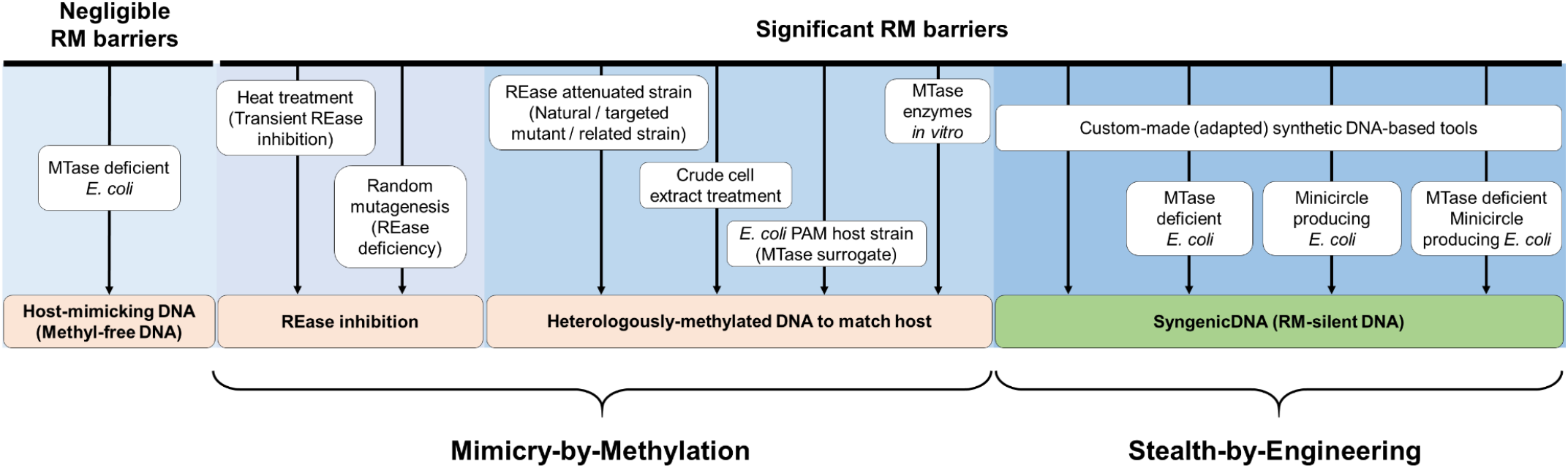
Approaches to overcome RM system-mediated genetic barriers in bacteria (adapted from^1^). Current approaches modify the methylation pattern of a genetic tool, either *in vitro* or *ex vivo,* to match that of the desired host to achieve mimicry by methylation. In contrast, SyngenicDNA methods evade RM systems by eliminating their target recognition sequences from DNA to create minimalistic RM-silent genetic tools, and achieve stealth-by-engineering during transformation.

**Supplementary Figure 2.**
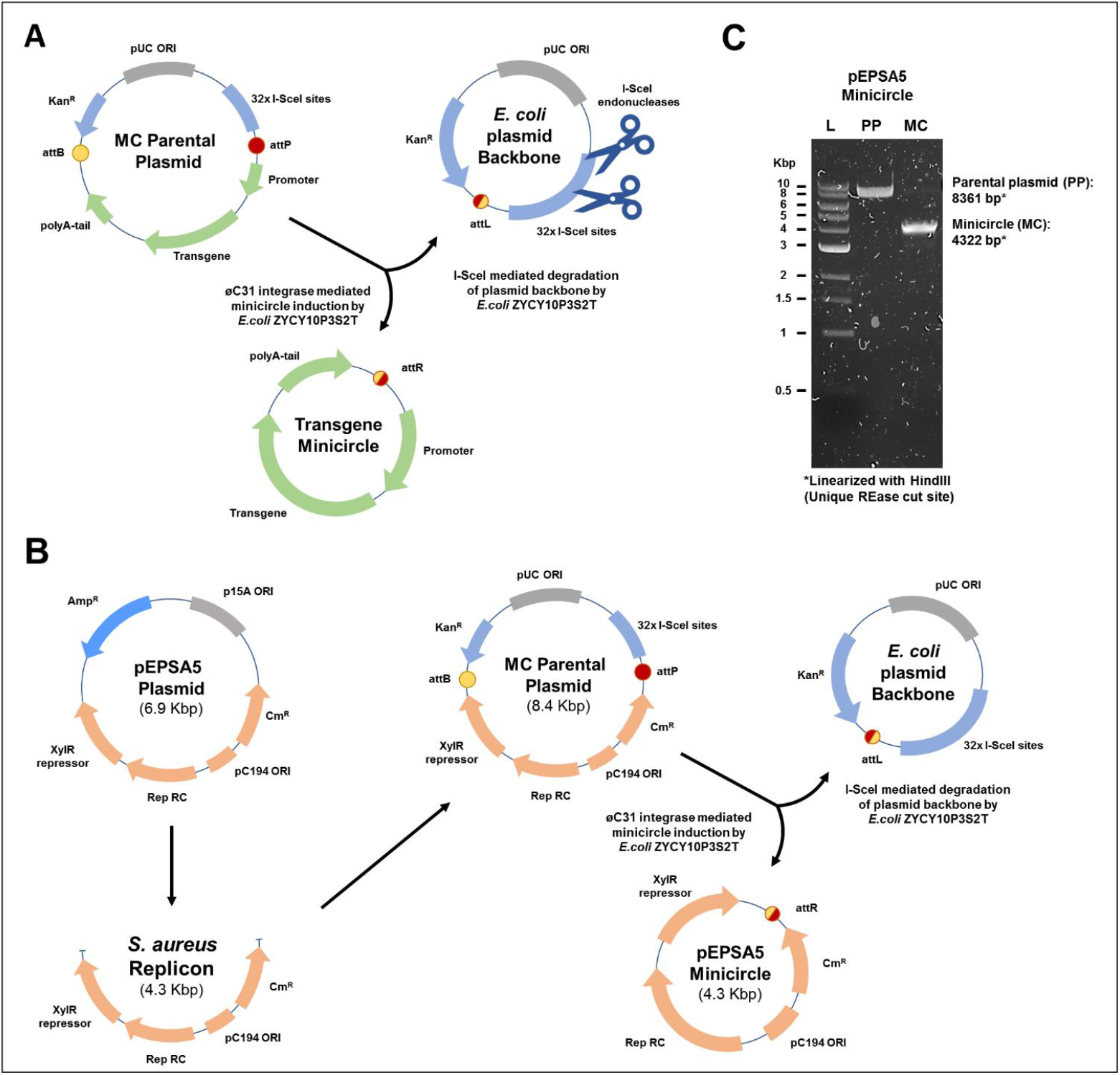
Repurposing of minicircle (MC) technology to produce minimalistic genetic tools for application in bacteria. **(A)** Current MC strategies (Kay *et al)* are applied to produce small circular expression cassettes for stable transgene expression in eukaryote hosts. Typically, a transgene cassette containing a eukaryote promoter, transgene, and polyA tail (green arrows) is attached to an *E. coli* plasmid backbone within a multiple cloning site flanked by attB and attP sites (bacterial and phage attachment recognition sites of the øC31 integrase enzyme, red and yellow circles respectively) to form a parental plasmid. The *E. coli* backbone also contains the antibiotic-selection marker Kan^R^ (blue arrow), a pUC origin (grey box) for high-copy-number autonomous replication in *E. coli,* and 32x tandem repeats of the I-SceI homing endonuclease recognition site (blue box) for I-SceI targeted degradation after MC induction. The øC31 integrase and I-SceI enzymes are arabinose inducible and encoded on the chromosome of *E. coli* ZYCY10P3S2T^2^. **(B)** In our repurposed bacterial MC strategy, a functional bacterial replicon/genetic tool takes the place of the eukaryotic transgene cassette. This allows for high-yield production of minimalistic genetic tools, which lack an *E. coli* replicon, for application in bacteria other than *E. coli.* We used the *S. aureus* replicon of the pEPSA5 plasmid to form a pEPSA5 minicircle that is 38% smaller than pEPSA5. **(C)** Restriction enzyme digestion of pEPSA5 parental plasmid (PP) and pEPSA5 minicircle (MC) following isolation from *E. coli* MC (ZYCY10P3S2T, a minicircle-producing strain). Plasmid DNA (500 ng), isolated prior to arabinose induction (PP) or 4-hours post induction (MC), was linearized with 1U of the unique cutter HindIII for 1 hour and resolved on a 1% agarose gel. Lane M, marker DNA (1 kb Ladder; NEB); lane PP, uninduced pEPSA5PP; lane MC, induced pEPSA5MC.

**Supplementary Figure 3.**
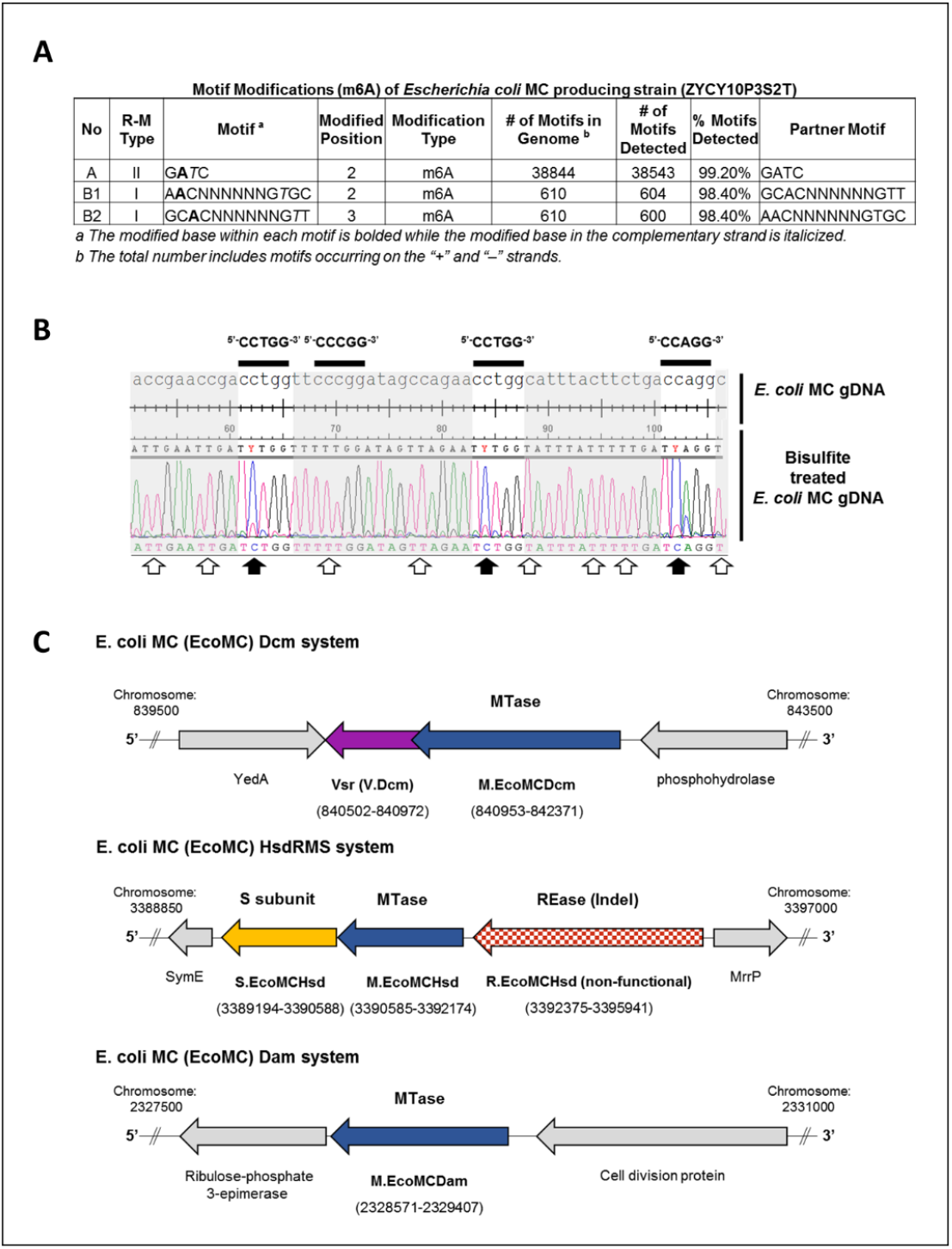
Methylation signatures present on *E. coli* MC (ZYCY10P3S2T) genomic DNA and the organization of responsible MTase gene clusters. **(A)** Detailed summary of 6-methyladenine (m6A)-modified motifs across the genome of *E. coli* MC (ZYCY10P3S2T, a minicircle-producing strain) detected by SMRTseq and Basemod analysis (the PacBio DNA modification sequence analysis pipeline, http://www.pacb.com/). RM systems were designated as Type I or II based on gene characterization through REBASE. ^*a*^The modified base within each motif is bolded while the modified base in the complementary strand is italicized. ^*b*^The total number includes motifs occurring on the + and – strands. **(B)** Summary of 5-methylcytosine (m5C) CCWGG-modified motifs on the *E. coli* MC genome. Sequence comparison and alignment of *E. coli* MC genomic region before and after bisulfite conversion. Unmethylated cytosine residues converted to thymine during bisulfite treatment are indicated by white arrows; m5C methylated cytosines protected from deamination are indicated by black arrows (present within CCWGG motifs, where W=A or T, but not CCCGG motifs). **(C)** A schematic representation showing the structure and genomic context of *E. coli* MC RM systems and orphan MTases. Gene assignments, nomenclature and genome coordinates publicly available at REBASE.

**Supplementary Figure 4.**
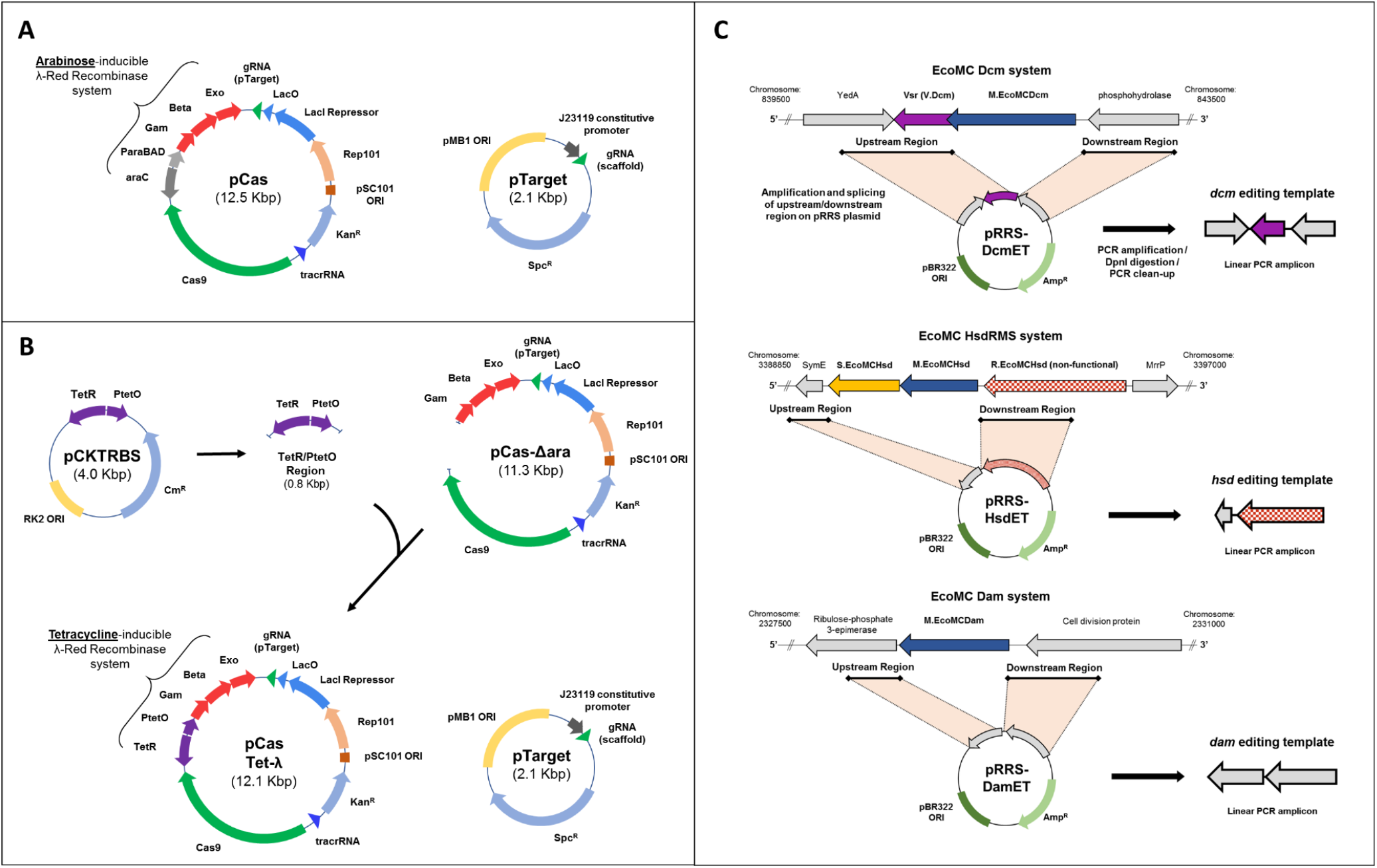
Engineering of an anhydrotetracycline-inducible CRISPR-Cas9/λ-Red recombineering strategy for scarless deletion of MTase genes within *E. coli* MC (ZYCY10P3S2T). **(A)** The original dual plasmid (pCas and pTarget) CRISPR-Cas9/λ-Red system developed by Jiang *et al.*^3^, with an arabinose inducible regulatory promoter/repressor module (araC-Pbad) controlling the λ-Red system (Gam, Beta, Exo). **(B)** Construction of the pCasTet-λ plasmid, a modified version of pCas. An 818-bp tetracycline-inducible regulatory promoter/repressor unit, TetR/PtetO, was amplified from pCKTRBS and spliced to a linear amplicon of pCas lacking the araC-Pbad module. The resultant plasmid, pCasTet-λ, contains λ-Red genes under transcriptional control of the TetR/PtetO regulatory cassette and can be used in combination with the original pTarget. **(C)** Assembly of DNA editing templates for MTase gene recombineering in *E. coli* MC. Approximately 400-bp regions from 5’ and 3’ of each MTase gene were spliced together onto a pRRS plasmid backbone to form the MTase deletion template plasmids (pRRSDcmET, pRRSHsdET, and pRRSDamET; where ET is editing template). These plasmids were used to amplify each MTase editing template prior to λ-Red recombineering.

**Supplementary Figure 5.**
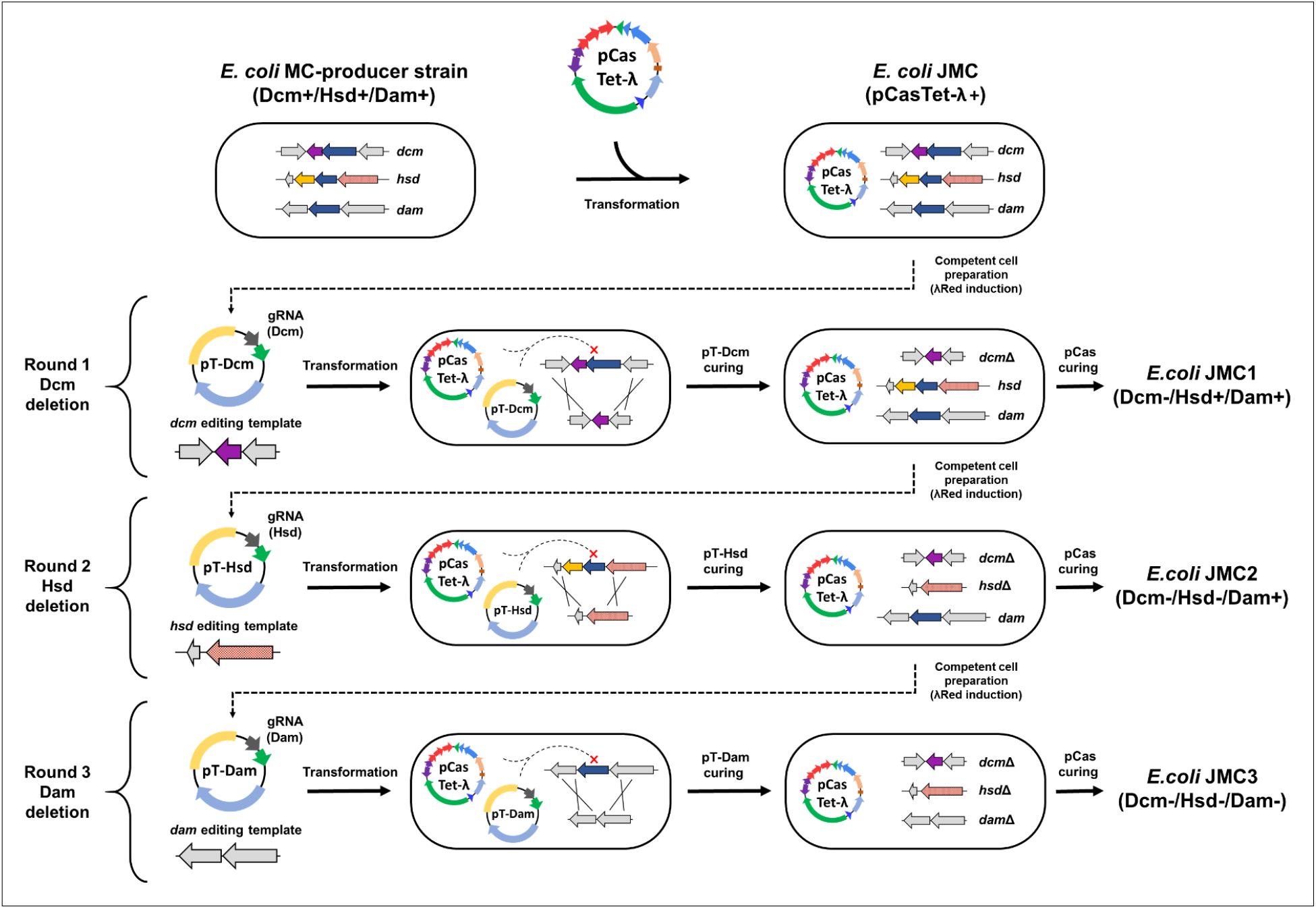
The CRISPR-Cas9/λ-Red recombineering scheme used in *E. coli* MC (ZYCY10P3S2T) for scarless MTase gene deletion. pTarget plasmids (pT-Dcm and pT-Hsd) each encode constitutively expressed gRNAs for Cas9-mediated targeting of MTase genes in unsuccessfully edited cells. gRNA sequences used are included in Supplementary Table 3.

**Supplementary Figure 6.**
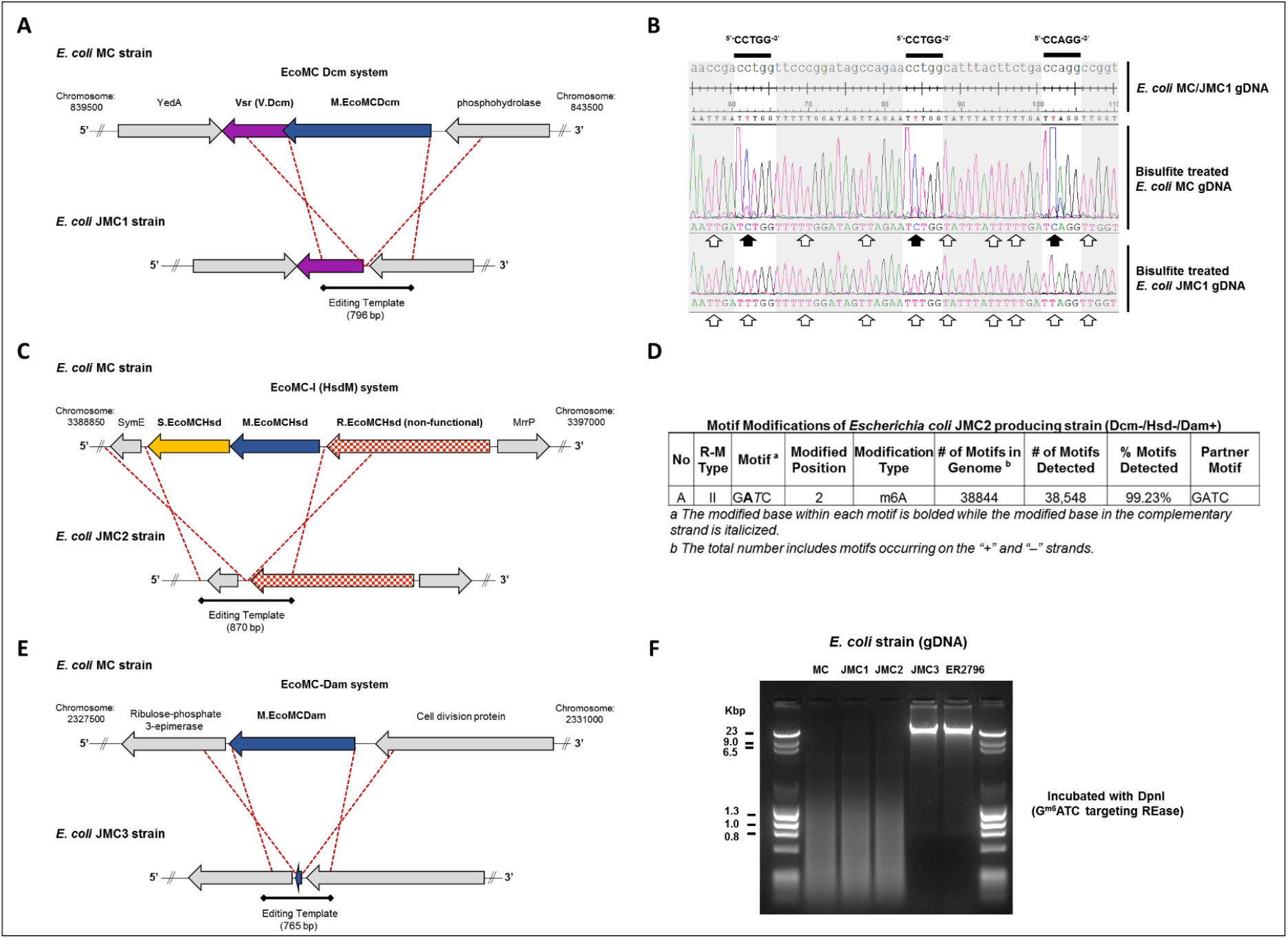
Schematic representations showing the context of genome editing in *E. coli* JMC-series strains along with phenotypic confirmation of MTase deficiencies. **(A)** Sequence confirmed *dcm* deletion in *E. coli* JMC1. **(B)** Comparison of Dcm activity in *E. coli* MC and *E. coli* JMC1 strains. Alignment of genomic regions before and after bisulfite conversion, highlighting the absence of m5C-modified CCWGG motifs on *E. coli* JMC1 gDNA (where W is A or T). White arrows indicate unmethylated cytosine residues converted to thymine during bisulfite treatment. Black arrows indicate m5C methylated cytosines protected from deamination. **(C)** Sequence confirmed *hsd* deletion in *E. coli* JMC2. **(D)** SMRTseq/Basemod summary of modified m6A motifs across the *E. coli* JMC2 genome, demonstrating the absence of methylated HsdS motifs (compared to the *E. coli* MC strain shown in **Supplementary Figure 3A**). **(E)** Sequence confirmed *dam* deletion in *E. coli* JMC3. **(F)** DpnI restriction of gDNA isolated from *E. coli* strains MC, JMC1, JMC2 and JMC3. Genoic DNA from the methyl-deficient E.coli ER2796 (NEB) is included as control. DpnI is a methyl-directed endonuclease that requires Gm6ATC for activity. JMC3 gDNA is resistant to DpnI cleavage indicating it is unmethylated at Dam (GATC) sites.

**Supplementary Figure 7.**
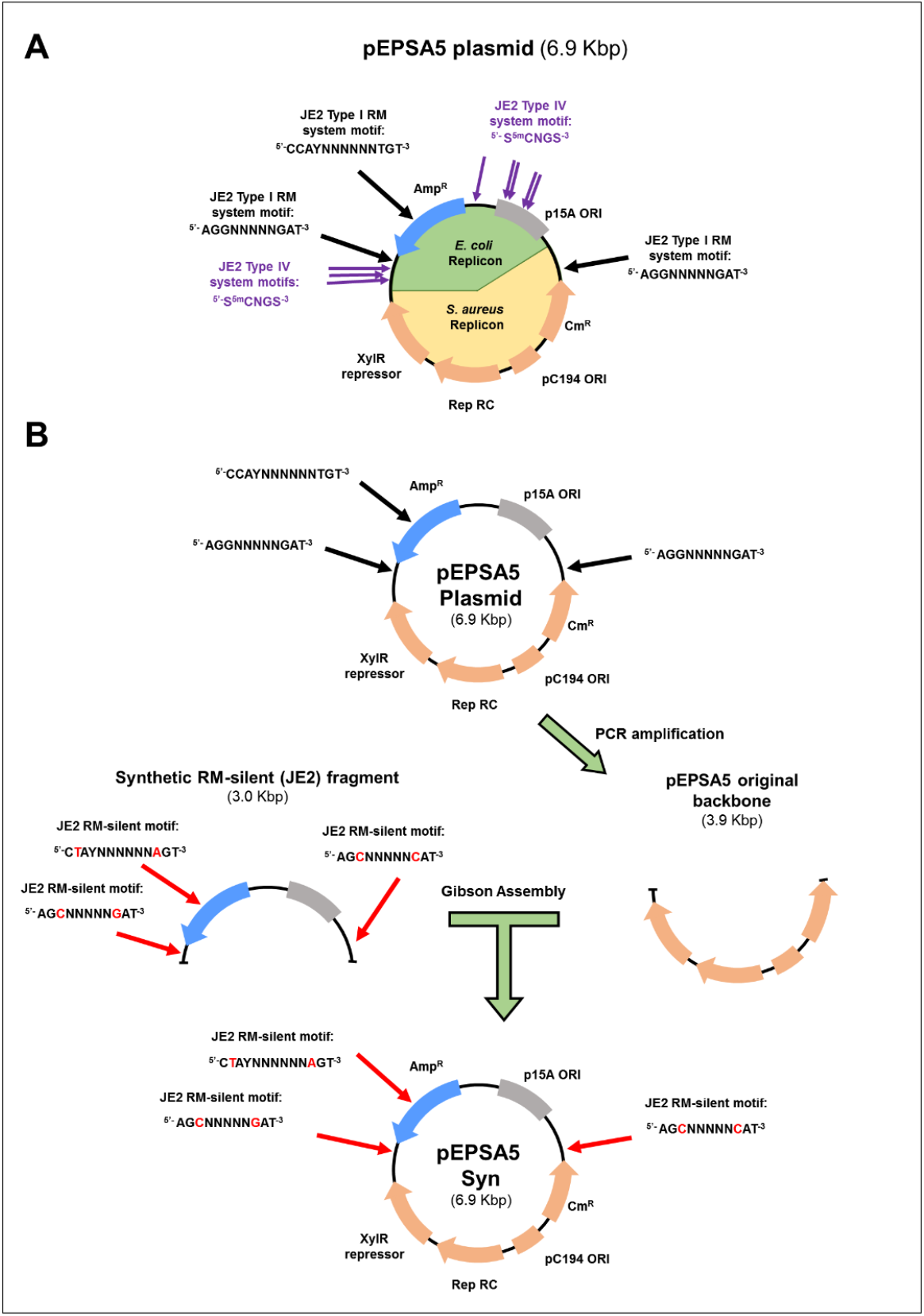
Schematic of pEPSA5 plasmid with *S. aureus* JE2 RM targets and construction of pEPSA5SynJE2. **(A)** Forsyth and colleagues described the original pEPSA5 *S. aureus–E. coli* shuttle vector^4^ This plasmid contains 11 individual *S. aureus* JE2 RM target motifs (Type I; n=3, and Type IV; n=8) that will be recognized and targeted for degradation upon transformation. **(B)** We assembled pEPSA5SynJE2 by replacing a 3-kbp fragment of pEPSA5 that contained three JE2 RM target motifs with a *de novo* synthesized RM-silent fragment. Black arrows indicate JE2 RM target motifs. Red arrows indicate those modified sites on the RM-silent fragment. Red letters indicate modified nucleotides. Type IV system targets are not shown, as these can be eliminated by propagation in a Dcm-deficient *E. coli* host. Both plasmids are 6850 bp in length and differed by only six nucleotides (99.91% nucleotide identity).

**Supplementary Figure 8.**
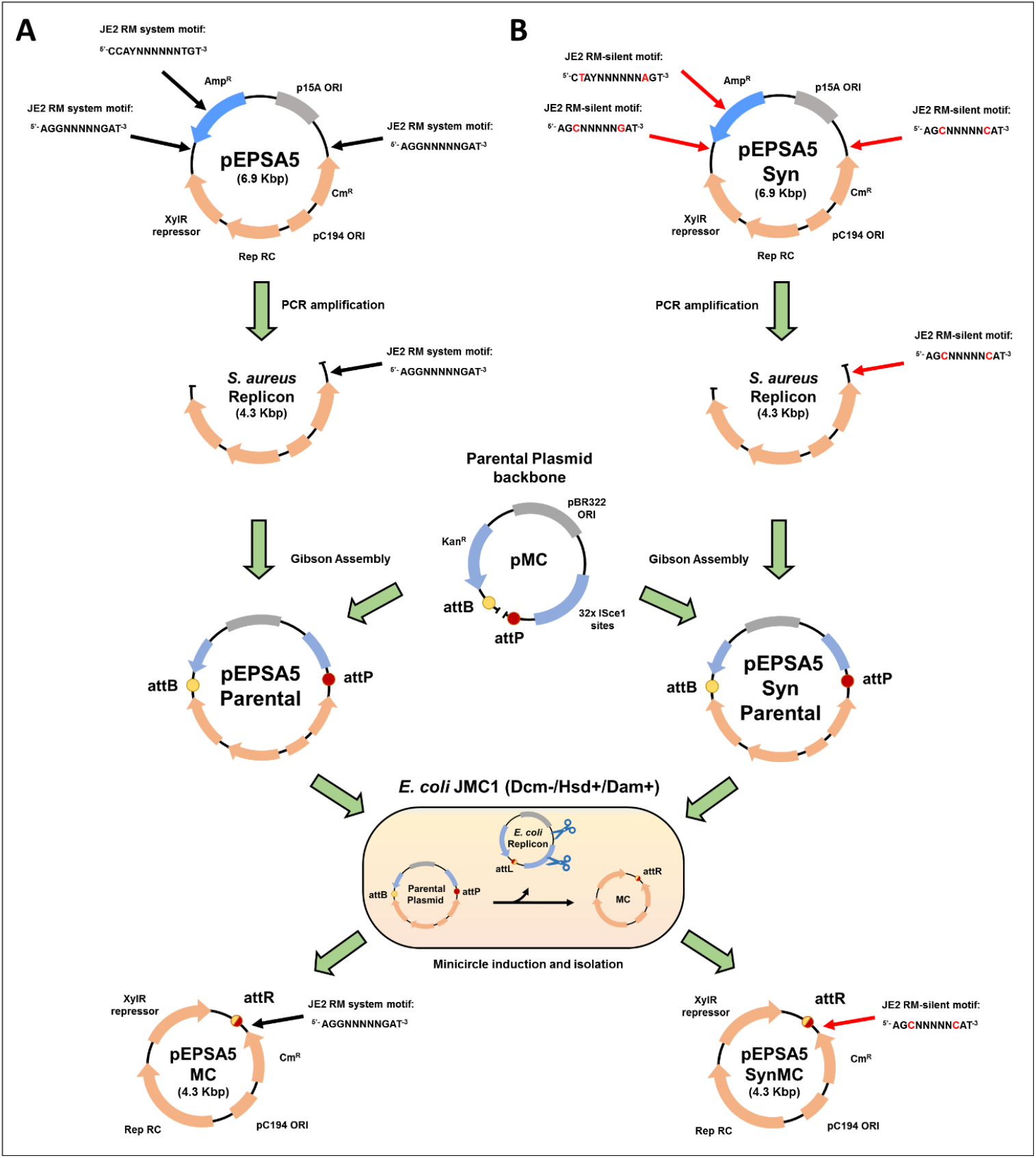
Assembly and propagation of pEPSA5- and pEPSA5Syn-based minicircles in *E. coli* JMC1. **(A)** The *S. aureus* functional replicon of pEPSA5, containing a single JE2 RM system target, was amplified to remove the original *E. coli* replicon. The *S. aureus* replicon was spliced to the pMC plasmid to form the pEPSA5 parental plasmid, which was transformed into competent *E. coli* JMC1 cells followed by arabinose induction of MC assembly. pEPSA5MC has a single JE2 RM system target. **(B)** This process was repeated for pEPSA5SynJE2, which is RM-silent with respect to JE2. pEPSA5MC and pEPSA5SynJE2MC plasmids differ by only the two nucleotides shown in red letters.

**Supplementary Figure 9.**
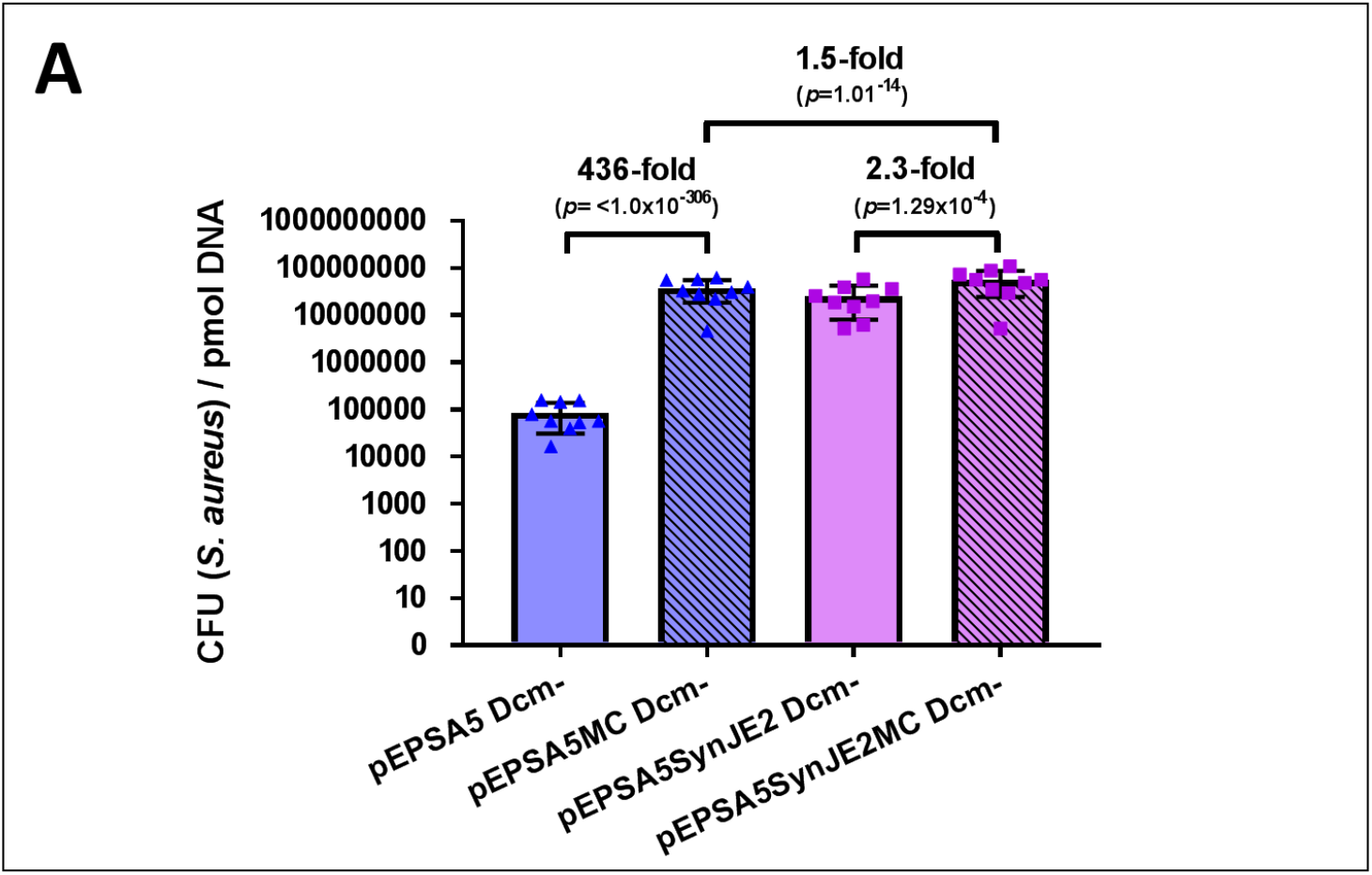
Secondary analysis of SyngenicDNA and pEPSA5-based SyMPL plasmid transformation efficiencies in CFU/pmol DNA. Data are means + SEM from nine independent experiments (three biological replicates with three technical replicates each).

## SUPPLEMENTARY TABLES

**Supplementary Table 1:**
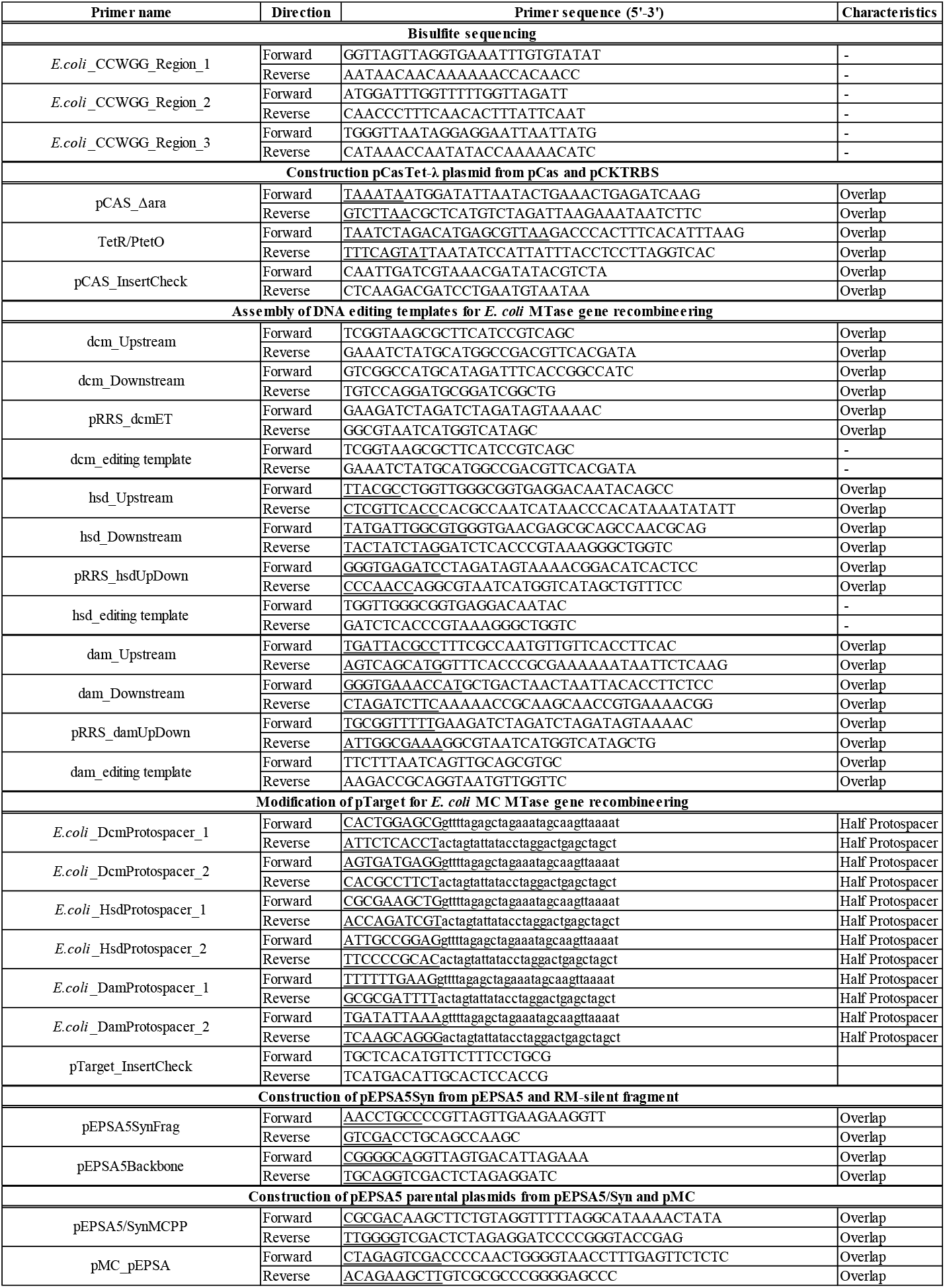
Oligonucleotides used in this study

**Supplementary Table 2:**
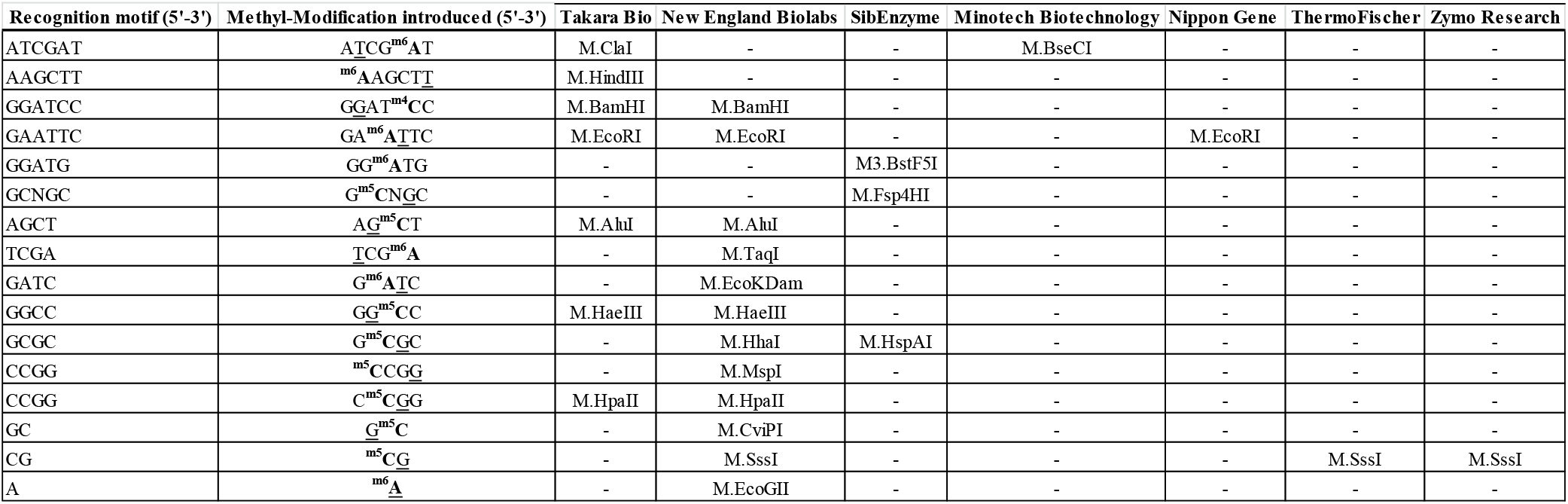
Methyltransferase enzymes commercially available for application in mimicry-by-methylation approaches.

**Supplementary Table 3:** *Staphylococcus aureus* JE2 colony counts for pEPSA5-based SyngenicDNA and SyMPL approaches

| Experiment 1: SyngenicDNA method |  |  |  |  |  |  |  |
| --- | --- | --- | --- | --- | --- | --- | --- |
| Competent cell Preparation | Experiment |  | CFU/μg plasmid DNA |  |  |  |  |
|  |  |  | pEPSA5 (Dcm +) | pEPSA5 (Dcm -) | pEPSA5SynJE2 (Dcm +) | pEPSA5SynJE2 (Dcm -) |  |
| OD<sub>600nm</sub> 0.86 | Biological Replicate 1 | Independant Replicate A | 0 | 385 | 10 | 159487.5 |  |
|  |  | Independant Replicate B | 10 | 532.5 | 32.5 | 264400 |  |
|  |  | Independant Replicate C | 0 | 505 | 42.5 | 219400 |  |
| OD<sub>600nm</sub> 0.80 | Biological Replicate 2 | Independant Replicate A | 15 | 757.5 | 30 | 210160 |  |
|  |  | Independant Replicate B | 2.5 | 655 | 47.5 | 212275 |  |
|  |  | Independant Replicate C | 2.5 | 795 | 42.5 | 228025 |  |
| OD<sub>600nm</sub> 0.93 | Biological Replicate 3 | Independant Replicate A | 10 | 2175 | 247.5 | 1077070 |  |
|  |  | Independant Replicate B | 10 | 2135 | 265 | 1268995 |  |
|  |  | Independant Replicate C | 12.5 | 775 | 105 | 663390 |  |
| Experiment 2: SyngenicDNA Minicircle Plasmid (SyMPL) method |  |  |  |  |  |  |  |
| Competent cell Preparation | Experiment |  | CFU/μg plasmid DNA |  |  |  |  |
|  |  |  | pEPSA5 (Dcm +) | pEPSA5 (Dcm -) | pEPSA5MC (Dcm -) | pEPSA5SynJE2 (Dcm -) | pEPSA5SynJE2MC (Dcm -) |
| OD<sub>600nm</sub> 1.67 | Biological Replicate 1 | Independant Replicate A | 0 | 13000 | 19780000 | 3430000 | 26190000 |
|  |  | Independant Replicate B | 0 | 9000 | 11050000 | 4230000 | 12470000 |
|  |  | Independant Replicate C | 0 | 3750 | 1630000 | 1180000 | 1840000 |
| OD<sub>600nm</sub> 1.56 | Biological Replicate 2 | Independant Replicate A | 185 | 35550 | 22150000 | 12930000 | 38980000 |
|  |  | Independant Replicate B | 135 | 11950 | 7960000 | 4460000 | 10360000 |
|  |  | Independant Replicate C | 185 | 17850 | 9920000 | 8780000 | 17220000 |
| OD<sub>600nm</sub> 1.52 | Biological Replicate 3 | Independant Replicate A | 535 | 32600 | 11840000 | 8140000 | 20020000 |
|  |  | Independant Replicate B | 385 | 12950 | 14260000 | 5760000 | 20100000 |
|  |  | Independant Replicate C | 295 | 35250 | 20380000 | 1380000 | 31260000 |

**Supplementary Table 4:** Fold changes in transformation efficiencies (CFU/μg) between pEPSA5 plasmid variants

| **Experiment 1: SyngenicDNA method** |  |  |  |  |
| --- | --- | --- | --- | --- |
| **Plasmids compared** | **Average -fold difference in counts** |  |  |  |
|  | **Fold difference** | **95% LB<sup>a</sup>** | **95% UB<sup>b</sup>** | **p-value** |
| pEPSA5 (Dcm +) versus pEPSA5SynJE2 (Dcm +) | 13.2 | 6.6 | 26.3 | $2.8 \times 10^{-13}$ |
| pEPSA5 (Dcm -) versus pEPSA5SynJE2 (Dcm -) | 493.8 | 399.3 | 610.5 | $<3.2 \times 10^{-308}$ * |
| pEPSA5 (Dcm +) versus pEPSA5 (Dcm -) | 139.4 | 80.5 | 241.7 | $2.5 \times 10^{-69}$ |
| pEPSA5SynJE2 (Dcm +) versus pEPSA5SynJE2 (Dcm -) | 5231.9 | 4494.7 | 6089.9 | $<3.2 \times 10^{-308}$ * |
| pEPSA5 (Dcm +) versus pEPSA5SynJE2 (Dcm -) | 68851.2 | 38393.2 | 123472.2 | $7.8 \times 10^{-306}$ |
| *a* and *b*: LB and UB are lower bound and upper bound of the 95% confidence interval |  |  |  |  |
| * p-value represented as an inequality as Stata software does not calculate p-values lower than this value |  |  |  |  |
| **Experiment 2: SyngenicDNA Minicircle Plasmid (SyMPL) method** |  |  |  |  |
| **Plasmids compared** | **Average -fold difference in counts** |  |  |  |
|  | **Fold difference** | **95% LB<sup>a</sup>** | **95% UB<sup>b</sup>** | **p-value** |
| pEPSA5 (Dcm -) versus pEPSA5SynJE2 (Dcm -) | 292.6 | 190.3 | 449.8 | $1.4 \times 10^{-147}$ |
| pEPSA5MC (Dcm -) versus pEPSA5SynJE2MC (Dcm -) | 1.5 | 1.4 | 1.7 | $1.0 \times 10^{-14}$ |
| pEPSA5 (Dcm -) versus pEPSA5MC (Dcm -) | 692.1 | 508.4 | 942.2 | $<3.2 \times 10^{-308}$ * |
| pEPSA5SynJE2 (Dcm -) versus pEPSA5SynJE2MC (Dcm -) | 3.5 | 2.3 | 5.4 | $1.8 \times 10^{-9}$ |
| pEPSA5 (Dcm -) versus pEPSA5SynJE2MC (Dcm -) | 1038.0 | 810.9 | 1328.7 | $<3.2 \times 10^{-308}$ * |
| *a* and *b*: LB and UB are lower bound and upper bound of the 95% confidence interval |  |  |  |  |
| * p-value represented as an inequality as Stata software does not calculate p-values lower than this value |  |  |  |  |

**Supplementary Table 5:**
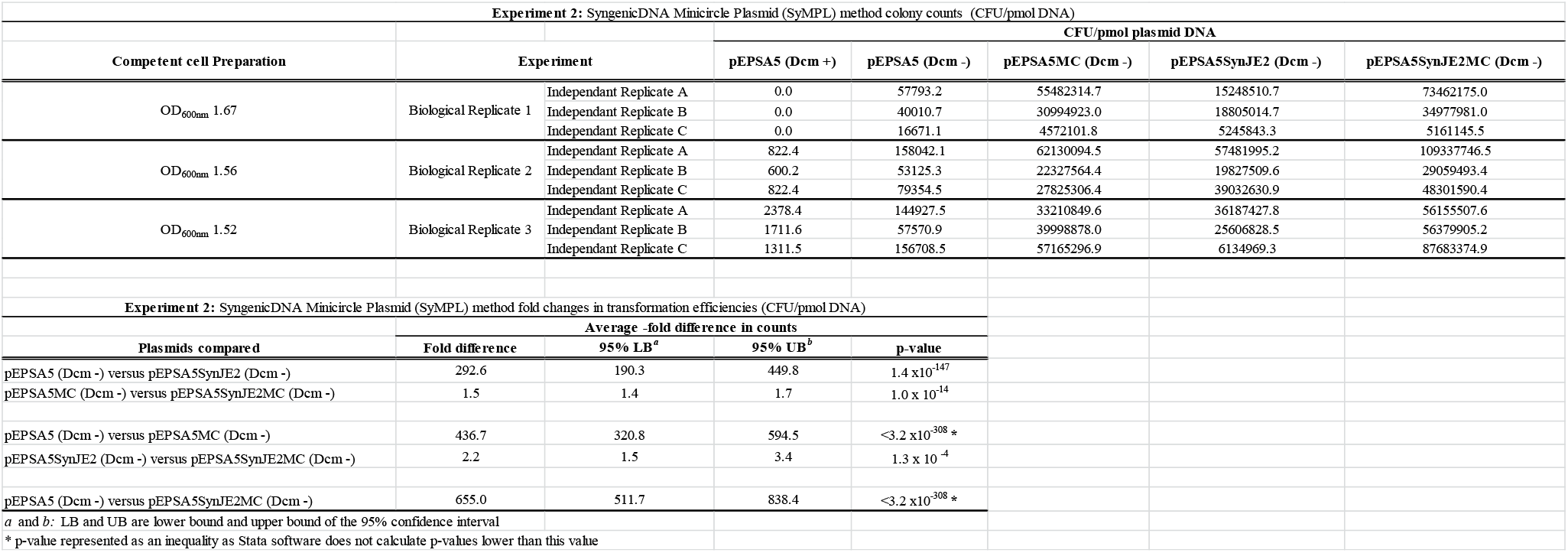
*Staphylococcus aureus* JE2 colony counts and fold changes in transformation efficiencies in CFU/pmol.

